# An affinity for brainstem microglia in pediatric high grade gliomas of brainstem origin

**DOI:** 10.1101/2022.06.15.496357

**Authors:** Liat Peretz Zats, Labiba Ahmad, Natania Casden, Meelim J. Lee, Vitali Belzer, Orit Adato, Shaked Bar Cohen, Seung-Hyun B Ko, Mariella G Filbin, Ron Unger, Douglas A Lauffenburger, Rosalind A Segal, Oded Behar

## Abstract

**Background:** High-grade gliomas (HGG) in children have a devastating prognosis and occur in a remarkable spatiotemporal pattern. Diffuse midline gliomas (DMG), including diffuse intrinsic pontine gliomas (DIPG), typically occur in mid-childhood, while cortical HGGs are more frequent in older children and adults. The mechanisms behind this pattern are not clear.

**Methods:** We used mouse organotypic slice cultures and glial cell cultures to test the impact of the microenvironment on human DIPG cells. Comparing the expression between brainstem and cortical microglia identified differentially expressed secreted proteins. The impact of some of these proteins on DIPGs was tested.

**Results:** DIPGs, pediatric HGGs of brainstem origin, survive and divide more in organotypic slice cultures originating in the brainstem as compared to the cortex. Moreover, brainstem microglia are better able to support tumors of brainstem origin. A comparison between the two microglial populations revealed differentially expressed genes. One such gene, Interleukin-33 (IL33), is highly expressed in the pons of young mice and its DIPG receptor is upregulated in this context. Consistent with this observation, the expression levels of IL33 and its receptor, IL1RL1, are higher in DIPG biopsies compared to low grade cortical gliomas. Furthermore, IL33 can enhance proliferation and clonability of HGGs of brainstem origin, while blocking IL33 in brainstem organotypic slice cultures reduced the proliferation of these tumor cells.

**Conclusions:** Crosstalk between DIPGs and the brainstem microenvironment, in particular microglia, through IL33 and other secreted factors, modulates spatiotemporal patterning of this HGG and could prove to be an important future therapeutic target.

**Key Points:**

- DIPGs divide more in the brainstem environment compared to the cortex.
- IL33 expression in brainstem and brainstem microglia is higher than in the cortex of young mice.
- IL33 propagates DIPG tumorigenesis, and its inhibition decreases DIPG proliferation.

**Importance of the Study:** Pediatric gliomas can arise in the supratentorial cerebral cortex with a median diagnosis age of 13 years, while diffuse midline gliomas (DMGs), present a median diagnosis age of 7 years, and appear most frequently in the pons. DMGs have a devastating prognosis and are currently untreatable. We tested the contribution of the brainstem microenvironment to the observed spatiotemporal pattern of pediatric HGGs. Our results support a model in which the brainstem microenvironment, particularly microglia, offers more supportive conditions for DIPG cells. By using the differences in gene expression between brainstem and cortical microglia we were able to identify factors that may contribute to the DIPG preference toward the brainstem. In this study we focused on the factor IL33, and provided evidence to support its contribution to the propagation of DIPGs in this specific anatomical location.

## Introduction

High grade gliomas (HGGs) constitute 8-10% of brain tumors in children and usually cause death within two years of diagnosis^1^. Pediatric gliomas can arise in the supratentorial cerebral cortex or in infratentorial regions, such as the cerebellum, brainstem, midbrain, thalamus, and spine^2^. Approximately half of pHGGs arise in midline locations, specifically in the thalamus and the pons, and are classified as diffuse midline gliomas (DMGs; pontine DMG is also known as diffuse intrinsic pontine glioma (DIPG)). While pontine and thalamic DMGs typically occur in mid-childhood, pediatric cortical gliomas occur in older children and young adults, and HGGs of later adulthood occur mainly in the frontotemporal lobes^2^.

Recent studies make it increasingly apparent that supratentorial and infratentorial pHGGs differ from one another and from their adult counterparts. In particular, the predominant oncogenic mutations vary depending on location and on the patient’s age. A striking example is the lysine to methionine mutation on residue 27 in histones H3.1 (H3.1K27M) and H3.3 (H3.3K27M) observed in approximately 80% of pontine DMG cases in younger children, while tumors with H3.3G34R/V mutations usually occur in the cerebral hemispheres of adolescents and young adults ^2–4^.

One possible explanation for this remarkable spatiotemporal pattern of DMGs may be a local advantage of the microenvironment of the infratentorial regions in a restricted time frame that fosters the development of tumors carrying these specific mutations. This assumption is consistent with the limited ability to induce tumorigenic transformations in older mice when using genes associated with DIPG mutations^5^.

A preference of pontine DMGs towards the young brainstem would imply that specific cells at a restricted time point in this location may be more suitable for the formation, progression, and survival of such tumors. If specific cell types in the brainstem indeed give an advantage to the formation of DIPG tumors, such cells should have differential anatomical and developmental properties. Consistent with this idea, accumulating evidence indicates that various cells in the CNS such as astrocytes, oligodendrocytes, and microglia show some anatomical and functional differences ^6–10^.

Evidence for interaction and cross talk between microglia, astrocytes, and vascular cells in adult forms of glioma have been previously demonstrated^11–13^. Pediatric brain tumors were also recently shown to be influenced by their environment ^14,15^. In this case, activated neurons in the micro-environment were shown to promote glioma growth.

In this work, we tested the possibility that pontine DMGs of brainstem origin are more attuned to brainstem surroundings, and subsequently aimed to find an anatomically specific DIPG-microenvironment crosstalk contributing to this compatibility.

## Materials and Methods

### Animals

C57BL/6 mice were obtained from ENVIGO (Rehovot, Israel). Animal handling adhered strictly to national and institutional guidelines for animal research and was approved by the Ethics Committee of the Hebrew University. Mice both males and females between day 2-4 after birth were used.

### Antibodies

A list of antibodies is in the supplementary methods.

### *Immunostaining and* Western blot

explanation is found in the supplementary methods.

### Patient-derived tumor cell cultures

All cultures were maintained as neurospheres in tumor stem medium (TSM) consisting of 50% Neurobasal(-A)/50% DMEM/F12, supplemented with B27™(-VitA) and growth factors as described previously ^14^ and in the supplementary methods.

### GFP+ or luciferase DIPG cell line preparation

Each patient derived HGG cell line was infected with Lenti-GFP/Luciferase viral supernatant and allowed to recover for 1 week. GFP-positive cells were isolated for purity by FACS (BD FACS Aria) and returned to culture. Luciferase positive cells were isolated via Puromycin (1μg/ml) selection. Validation of the sorting was done using FACS analysis and by observing the cell fluorescence under the microscope (with at least a 90% positive GFP rate).

### Slice Cultures preparations

Tumor cells (10,000 cells/µl) were re-suspended in Matrigel (Corning, NY, USA) and TSM (1:1) and injected (1µl) using Narishige MM-3 micromanipulator (Tokyo, Japan). The cells were systematically injected in parallel into both the brainstem and cortex of P2-4 mouse brains, two injections per region. The injections of each region were performed consecutively, alternating the order of injections between brainstem and cortex. After injections, the brain regions were gently separated and tissue pieces were incubated on top of 40µm cell filters. Each well was filled with medium (Neurobasal, 15mg/ml glucose, AmphoB 25 µg/ml, Pen-Strep, and B27 (without vitA)) up to the level of the filter to allow diffusion. Incubation was for 24-72h in 5% CO2, 37°C.

### Ex-vivo Multiphoton imaging and quantification

Each injected cortex and brainstem pair was incubated for 24hrs, then sealed with a cover slip (moisturized with PBS) and imaged live, using the large image option of *Nis-Elements*, and the maximal width in the Z stack option so as to capture all of the injected GFP+ cells. Analysis is described in the supplementary methods.

### *Ex-vivo* GFP quantification analysis

After 24h, slices were dissociated, and analyzed in a Fortessa LSR cell analyzer (BD Biosciences, CA, USA). We made sure to record all of the events from each sample (about 3-8 million events per sample), so that quantification represents the absolute GFP+ cell count per brain region injected. For each cell line, at least 9 brains were injected, in at least 3 injection sessions.

### *Ex-vivo* CellTrace proliferation analysis

Glioma cells were labeled with CellTrace reagent and injected into mouse brains as previously described (CellTrace™ Violet Cell Proliferation Kit, for flow cytometry, Thermofisher, MA, USA). A T0 measurement (maximal CellTrace intensity) was obtained per trial and used to determine the percentage of dividing cells after 24h or 72h incubation.

### Ex-vivo cell death analysis

Following slice culture incubation of either 6 or 24h, injected tissue samples were dissociated and stained with 100µg/ml Propidium Iodide (PI) (Thermofisher, MA, USA). Flow cytometry was used to quantify the percentage of PI positive cells out of GFP positive cells.

### Microglia and astrocytes cell culture

Preparation of mixed glia primary cell cultures and astrocytes from 2-3 day old mice was previously described^16^. Microglia were prepared as described here^17^. Briefly, to produce an astrocyte cell culture, the cells are plated in very low density, 20ng/ml EGF (PeproTech, Rehovot, Israel) is applied in the first few day, and later on the culture gets shaken overnight, treated with AraC (0.1mM, Sigma Aldrich) and passaged twice in order to ensure purity. For microglia isolation, mix glial cells were plated densely and treated with 0.5ng/ml gmCSF (PeproTech, Rehovot, Israel), until the cells are easily isolated from the culture and plated for experiments. An extended explanation is found in the supplementary methods.

### qPCR DNA quantification

Testing of cell number with DNA based qPCR and primers was carried out based on a previous paper with some modifications^18^. Briefly, DNA from a controlled number of glioma and mouse (astrocytes and microglia) cells was acquired, then a qPCR reaction (iTaq Universal SYBR Green Supermix, BioRad, Rishon Le Zion, Israel) using either a human specific primer or a primer designed to detect both human and mouse cells, was performed on the control DNA samples (Oligonucleotide sequences can be found in the supplementary methods). The results were used to create transformation formulas from cycle threshold (CT) to cell number/DNA amount with a high prediction power (R^2^=0.99 for both primers). To test the strength of these transformation formulas, sets of quantified human and mouse DNA samples (from 100% human DNA, 90% human with 10% mouse, etc., all the way to 100% mouse DNA) were tested. Using our transformation formula, we were able to accurately estimate the ratio of mouse and human cells in a mix culture, with a strong predictive value (R^2^=0.98).

### Co-culture proliferation experiments

8,000 human HGGs were placed on top of mouse glial cell cultures (12 well)-astrocytes and microglia, originating either in the brainstem or the cortex of the same mice. For each condition, within each trial, T0 wells were acquired about 2 hours after the trial began. The co-culture was then incubated for 72 hours in glioma medium, after which DNA was acquired using the Monarch DNA cleanup kit (NEB, Ipswich, MA, USA), and DNA-based qPCR quantification was done, as is described above.

### Microglial conditioned medium preparation

For each trial a confluent microglial cell culture was prepared, taking care to maintain similar conditions for the brainstem and cortical derived microglia. Once cell cultures reached similar confluence, the cells were thoroughly washed to remove any residual serum, and TSM without factors was added. For each 12well well, 1.5ml of medium was added. After 72hrs the medium was collected and filtered.

### luciferase based viability assay

Luciferase labeled DIPG cells were plated on top of purified microglia or astrocytes (670 cells/well), in a 96-well plate. Following a 72h incubation, luciferase activity was measured using LUMIstar® Omega (BMG LABTECH, Ortenberg, Germany) microplate luminescence reader. Each experiment consisted of a minimum of four wells per condition, and each experiment was repeated at least six time per cell line.

### Three dimensional growth experiments

Recombinant human IL33 (50ng/ml, PeproTech, Rehovot, Israel) was added to a TSM and Matrigel mixture (1:1 ratio) with DIPG cells highly concentrated to create very dense cell-matrigel droplets. These marked a specific starting point. Control droplets were also added. After initial solidifying of the gel, the radius of the droplet was expanded by adding more Matrigel-TSM mix. After final solidification, TSM based medium was added on top of the Matrigel drops to cover them. The diameter of each DIPG drop was measured after 2 hours and 7 days. ImageJ was used to measure the change in diameter. In each trial, at least 7 droplets for each condition were used.

### Proliferation experiments

CellTiter-Glo® Luminescent Cell Viability Assay (Promega, Madison, WI, USA) was used as described in the supplementary methods in the ‘Recombinant IL33 concentration calibration’.

### Colony count and average colony size experiments

400 DIPG cells were plated in 0.3cm^2^ wells. This low density plating allowed clear segregation of colonies. For each condition, at least 7 wells were plated per repeat. After 5 days incubation the cultures were analyzed using Incucyte® S3 (Sartorius, Göttingen, Germany) Live-Cell Analysis system (whole well image X4).

### RNA-Sequencing analysis

RNA data processing, data analysis, and subcellular localization analysis are described in the supplementary methods.

### PedcBiPortal Gene expression

Data and the clinical data associated with each sample were downloaded from the PedcBioPortal (https://pedcbioportal.kidsfirstdrc.org/)^19,20^ on February 2022. Data analysis is described in the supplementary section.

### mIL33 neutralization trial in *ex-vivo* Slice cultures

DIPG cells, labeled with CellTrace Dye, were injected with 1μg/ml of mIL33 antibody or isotype control antibody into each side of the cortex and of the brainstem, then slice cultures were prepared. After 3 days, the proliferation of DIPGs within each condition was measured using FACS analysis. An additional cortical injection was used to exclude any brains in which no brainstem preference was detected from the statistical analysis, excluded brains: DIPGXIII 3/13, DIPGXXV 1/12, BT869 3/14.

### shRNA IL1RL1 knockdown

was performed essentially as previously described^21^. For IL1RL1 knockdown experiments we used Mission® shRNA PLKO lentiviral vectors which include puromycin selection marker: TRCN0000358832/sh-IL1R-1, and TRCN0000058513/sh-IL1R-2. For a control we used an empty vector (pLKO.1-puro), all vectors purchased from Sigma Aldrich (St. Louis, MO, USA). Details are described in the supplementary section.

### qPCR Analysis

was performed essentially as previously described^21^. Differential expression was determined using the delta CT method. Each primer set was tested on human and mouse samples to make sure each primer is species specific. Further detail in supplementary methods.

### Statistical analysis

Generally, non-parametric, coupled (Wilcoxon) or un-coupled (Mann-Whitney), one tailed statistical tests were applied. Described in detail in the supplementary methods.

## Results

### Preference of pediatric HGGs of brainstem origin towards the brainstem

To test the intrinsic compatibility of DIPGs to their anatomical origin, we adopted an *ex-vivo* approach, which allowed us to test the initial response of DIPGs to their microenvironment. We injected equal numbers of GFP labeled patient-derived DIPG cells into the brainstem and cortex of the same brains followed by organotypic slice cultures. Using two photon microscopy (Fig. 1a-c), we detected higher numbers and a broader distribution of DIPG cells in the brainstem slices in comparison to the cortex.

**Figure 1:**
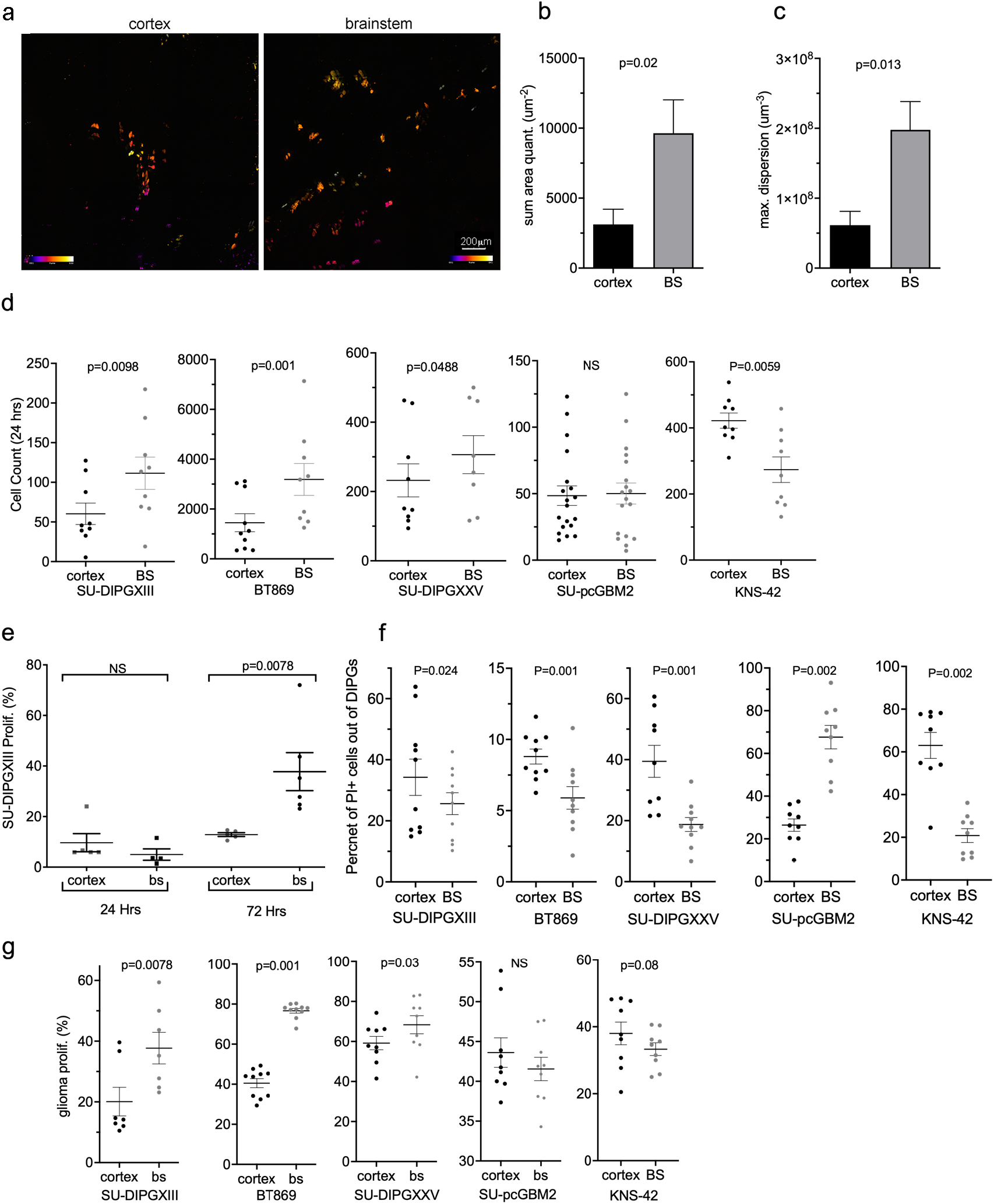
Preference of pediatric HGGs of brain stem origin towards the brainstem. (a) Two photon micrograph of infiltrating SU-DIPGXIII cells expressing eGFP, 24h post injection into P2 mouse brain. (b) Nis-elements analysis of eGFP positive cell amount in each micrograph (Mann-Whitney U test; n=6). (c) Diffusion quantification of eGFP positive cells within each micrograph measuring the three-dimensional maximal area (um^3^) (Mann-Whitney U test; n=6). (d) FACS analysis quantifying the total glioma count in each sample, 24hrs post injection ±SEM (Wilcoxon’s signed rank test, one tailed; SU-DIPGXIII n=9, BT869 n=10, SU-DIPGXXV n=9, SU-pcGBM2 n=9, KNS-42 n=9). (e) FACS analysis of CellTrace dye staining of GFP+ SU-DIPGXIII cells. Data presented as the average percent of proliferation of glioma population ±SEM (Wilcoxon’s signed rank test, one tailed; n=6-7). (f) FACs analysis of PI staining of GFP+ glioma cells 6hrs post injection into the cortex and brainstem of the same brain. Each point represents the percentage of PI+ cells out of the GFP+ population, indicating glioma cell death ±SEM. (Wilcoxon’s signed rank test, one tailed; -DIPGXIII n=10,BT869 n=10, SU-DIPGXXV n=10, SU-pcGBM2 n=9, KNS-42 n=9). (g) FACS analysis of CellTrace dye staining of GFP+ glioma cells 72h post injection into the cortex and brainstem of the same brain. Each point on the graph represents the percentage of GFP+ cells which underwent division (at least one), out of the entire GFP+ cell population ±SEM. (Wilcoxon’s signed rank test, one tailed; -DIPGXIII n=7,BT869 n=10, SU-DIPGXXV n=9, SU-pcGBM2 n=11, KNS-42 n=9).

To quantify the number of tumor cells in each region we used FACS analysis. In all DIPGs tested, more cells were present in the brainstem. In contrast, the pediatric HGG of cortical origin KNS-42, showed a strong preference toward the cortex, while another cortical line, SU-pcGBM2, showed no preference to either site (Fig. 1d and Fig S1).

To test proliferation, we used CellTrace™ dye at 24h and 72h (Fig. 1e). At 24h there was very little DIPG division overall, suggesting improved survival as opposed to proliferation in the brainstem at this time point. To test this directly, tumor cell death in the slice cultures was analyzed. After 6h (Fig. 1f), cells from all DIPG lines died less in the brainstem while SU-pcGBM2 showed an opposite response. At 24h (Fig S2) only BT869 and pcGBM maintained their original preference while the other lines showed no significant difference (Fig. S2). Surprisingly, KNS-42 died less in the brainstem. We then tested proliferation after 72h, when initial difference in proliferation was detected. We found significantly more DIPG division in the brainstem. In contrast, SU-pCGBM2 and KNS-42 showed a trend toward the cortex although this was not statistically significant (Fig 1g and Fig. S3).

### Brainstem microglia increase the accumulation of HGGs of brainstem origin

To assess the glial cells’ contribution, we tested the proliferation of SU-DIPGXIII on mixed glial cultures from the brainstem and cortex using a DNA based qPCR approach. We calculated the estimated proportion of DIPG cells according to cycle threshold, based on calibration curves prepared with known numbers of human and mouse cell mixes (Fig. S4). Interestingly, DIPG cells proliferated significantly more when incubated with brainstem glia (Fig 2a). Since microglia are known to infiltrate the tumor microenvironment and contribute considerably to glioma development^22–25 22–25^, we decided to test their role. DIPGs were plated with purified microglia from each brain region. Tumor cell number was estimated using the DNA based qPCR quantification described above. All HGGs of brainstem origin accumulated significantly more on brainstem microglia, while SU-pCGBM2 did not (Fig. 2b, Fig S5a). In contrast, DIPGs showed no anatomical preference on astrocytes (Fig. 2c, Fig S5b), while SU-pcGBM2 proliferated more on cortical astrocytes. Although microglia and astrocytes are altered and activated in culture, we assumed these alterations occur in cells from both regions, allowing an informative comparison.

**Figure 2:**
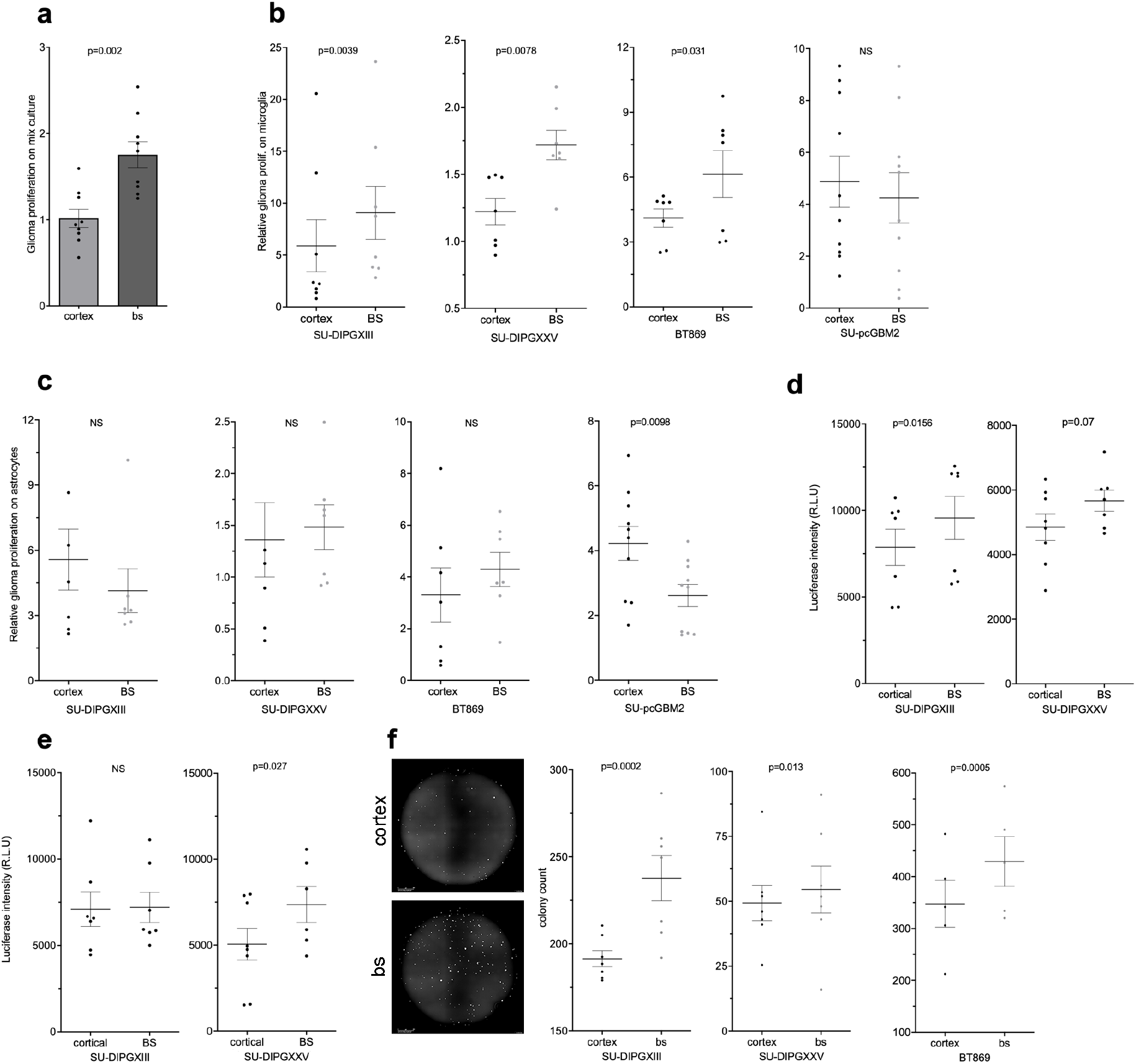
Microglia are responsible for brainstem differential effect on brainstem derived glioma cell lines (DIPGs). a) Glioma cell quantification using qPCR on human genomic DNA; DNA acquired from co-cultures of mouse mixed glia and human SU-DIPGXIII. The data is presented as the average fold change per trial, after 72h compared to T=0±SEM. Each point on the graph represents the average of at least 4 repeats. (Wilcoxon’s signed rank test; n=9. One tailed). (b+c) Glioma cell co-culture proliferation. The proportion of the tumor within the co-culture was estimated using qPCR on human/mouse genomic DNA; the DNA was acquired from mouse microglia (b) or astrocytes (c) and human glioma co-cultures after three days of incubation. The data is presented as the average fold change per trial after 72h, compared to T=0±SEM (each point on the graph represents an average of at least 3 repeats). (Wilcoxon’s signed rank test, one tailed; SU-DIPGXIII n=8, SU-DIPGXXV n=7, BT869 n=7, SU-pcGBM2 n=11). (d+e) DIPG co-culture proliferation quantified with a luciferase based assay. Luciferase positive DIPG lines were incubated on top of brainstem or cortical derived microglia (d) or astrocytes (e). After 72hr the luciferase intensity was quantified, and the data is presented in Relative Luminescence Units (each point on the graph represents the average of at least 3 repeats). (Wilcoxon’s signed rank test, one tailed; SU-DIPGXIII n=8, SU-DIPGXXV n=8). (f) The effect of microglial derived conditioned medium on clonability properties of DIPG cultures (representative image -left). The colony count per well is presented ±SEM, each point represents a median count of 8 repeats. (Fisher’s Combined Probability Test; SU-DIPGXIII n=7, SU-DIPGXXV n=5, BT869 n=7. Within each trial, a Mann Whitney U test was performed to obtain the P-values. One tailed).

To validate the DNA qPCR results, two DIPG lines labeled with luciferase were used to estimate DIPG accumulation on microglia or astrocytes, with similar results (Fig 2d, 2e, S5c). As a final validation step for the effects of microglial cells on DIPGs, we used anti-human specific antibody to identify and count SU-DIPGXIII cells with a comparable outcome (Fig. S6). Altogether, a clear intrinsic preference of DIPGs to brainstem microglia was documented.

To test the involvement of secreted factors, conditioned medium from brainstem and cortical microglial cell cultures (incubated without tumor cells) was used in a colony formation assay. In all DIPGs tested, the number of colonies was significantly higher with brainstem conditioned media (Fig. 2f). Thus, at least part of the propagative influence of brainstem microglia is mediated by secreted factors.

### Differential gene expression between brainstem and cortical microglia

Based on the observed functional difference between microglia of brainstem and cortical origin, we analyzed the differences in gene expression between them. Since HGGs and microglial cells might impact each other’s gene expression, microglia were incubated with SU-DIPGXIII. For gene expression analysis, the microglia-DIPGXIII sample data was aligned to a combined human and mouse reference genome. Though differences were detected, no statistically significant differences could be found in gene expression of the human genes (SU-DIPGXIII) by applying a uniform statistical cutoff. In contrast, mouse microglia revealed a significant number of differentially expressed genes (Fig. 3a, <u>Table S1</u>).

**Figure 3:**
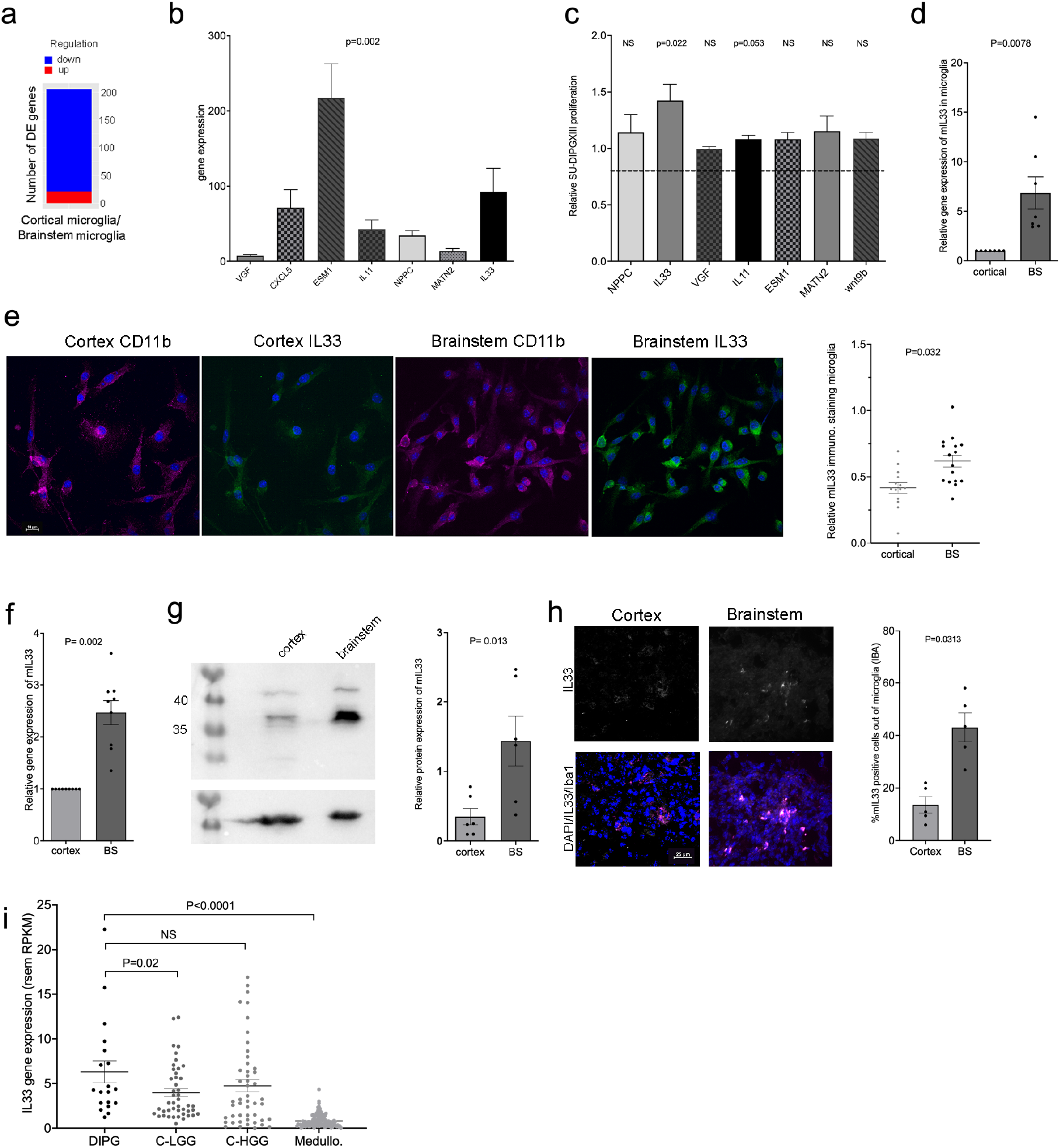
Differential gene expression between brainstem and cortical microglia. (a) Global gene analysis of brainstem and cortical microglia using RNA seq approach. Chart summarizing number of differentially expressed genes (DEGs) where DEGs are genes that had abs{log2FoldChange}>1 and padj<0.05, after DESeq2analysis. Regulation was determined by the sign of log2FoldChange, wherelog2FoldChange>0 indicated up-regulation while log2FoldChange<0 indicated down-regulation in the first sample of each comparison. The comparison shown refers to gene expression change of cortical microglia on SU-DIPGXIII against brainstem microglia on SU-DIPGXIII. There are 21 up-regulated and 185 down-regulated genes in this comparison (n=3 from each sample analyzed). (b) Quantitative PCR analysis was done on at least 6 separate SU-DIPGXIII + microglia co-culture trials, per gene tested. The values are expressed as the average fold change in brainstem co-cultures in comparison to the corresponding cortical co-cultures, ± SEM. (Wilcoxon’s signed rank test; NPPC and IL33 n=6; MATN2, VGF, IL11 and ESM1 n=9, wnt9b n=8. One tailed). (c) Screening of selected secreted factors, isolated from RNAseq analysis, for positive proliferation effects on SU-DIPGXIII. Glioma cells were incubated with control medium or with a factor for 72h, after which proliferation was measured using the CellTiterGlo luminescence system. The data is presented as the average fold change of the secreted factor’s effect as compared to the control ± SEM. (Wilcoxon’s signed rank test; NPPC, MATN2 n=14, IL33, VGF n=20, Epha3, Wnt9b, IL11, ESM1 n=9. One tailed, Bonferroni correction for multiple comparisons was applied). (d) Quantitative PCR analysis of mouse IL33 gene expression in pure microglial cell cultures. The values are expressed as the average fold change in brainstem cultures in comparison to the corresponding cortical cultures ± SEM. (Wilcoxon’s signed rank test; n=7). (e) Representative immunofluorescence for IL33 (green) and CD11b (magenta) on brainstem or cortical microglia (scale bar 10μm). Analysis was performed with Nis-elements by measuring the ratio of mIL33 positive cells out of CD11b positive cells. Quantification was done with the ‘sum area quantification’ tool. (Fisher’s Combined Probability Test; n=3.Within each trial, a Mann Whitney U test was performed to obtain the P-values, n=6 fields. One tailed). (f) Quantitative PCR analysis of mouse IL33 gene expression in the brainstem and cortex of P2 mice. The values are expressed as the average fold change in the brainstems in comparison to the corresponding cortexes, ± SEM. (Wilcoxon’s signed rank test; n=9) (g) A representative Western blot analysis of mIL33 expression in the brainstem and in the corresponding cortex of P2 mice. The graph shows a quantitative analysis of the Western blot trials, using GelQuant express, and then normalizing the mIL33 expression to the actin on the same membranes. At least 6 comparisons of brainstem-cortex were performed (Wilcoxon’s signed rank test; n=6). (h) Representative immunofluorescence for IL33 (white) and Iba1 (magenta) on a brain slice, from either the brainstem or the cortex. (scale bar 25μm). Analysis was performed with Nis-elements by measuring the percent of mIL33 positive cells out of Iba1 positive cells. Quantification was done with the ‘sum area quantification’ tool. (Wilcoxon’s signed rank test; n=5). (i) Human IL33 gene expression data obtained from PedcBioPortal. The gene’s expression within biopsies from DIPG samples was compared to cortical low and high grade gliomas and to medulloblastoma biopsies. Each point on the graph represents a single human sample. FPKM= fragments per kilobase of exon per million mapped fragments, RSEM= RNASeq by Expectation-Maximization (Mann Whitney U test was performed).

To select relevant genes for analysis, we used the Uniprot database, and identified 92 protein coding genes of secreted or membrane-associated proteins (<u>table S1</u>). We then focused on potentially novel candidates with a high fold change and known links to cancer, particularly glioma. Based on these criteria, we selected 7 genes. To validate their differential expression, RNA from microglia-SU-DIPGXIII cocultures was tested using mouse specific qPCR analysis (Fig. 3b).

Next, we tested the ability of proteins coded by these genes to induce the proliferation of SU-DIPGXIII (Fig. 3c). Out of the 7 candidates, only IL33 significantly enhanced HGG proliferation. To test whether IL33’s differential expression is intrinsic, we performed qPCR (Fig. 3d) and immunostaining (Fig. 3e) on purified microglia. Results indicated that this difference is indeed intrinsic. Consistent with this, the expression level of IL33 mRNA transcripts and protein is higher in the brainstem of mice at a young age (Fig3f, 3g), with this higher expression being at least partly due to the expression in brainstem microglia (Fig 3h, Fig S7). Furthermore, according to qPCR analysis, this anatomically differential expression significantly lessens at the age of two months and is most significant from birth up to 8 weeks (Fig. S8)

Interestingly, IL33 expression in DIPG patients is significantly higher than in patients with cortical low grade glioma or medulloblastoma (Fig 3i). The higher expression levels of IL33 in human DIPGs and pediatric cortical HGG samples is consistent with the IL33 data in adult HGGs^26–28^. Since cellular resolution is lacking, the source of IL33 in DIPGs may be the micro-environment, as our data suggests, while in the cortical HGGs the source could be the glioma cells, as has been shown for adult HGGs^26,27^.

### IL33 is able to impact DIPG tumor cells

To find the working concentration of IL33, we carried out a dose-response proliferation assay using DIPGXIII (Fig. S9), and selected a concentration of 50ng/ml, which is consistent with previous studies^29–31^. We then tested the effect of recombinant IL33 on DIPG proliferation (Fig. 4a), 3D growth (Fig. 4b and Fig. S10), and colony formation (Fig. 4c, Sup. Fig. S11). In all DIPG lines tested, IL33 significantly enhanced tumorigenesis.

**Figure 4:**
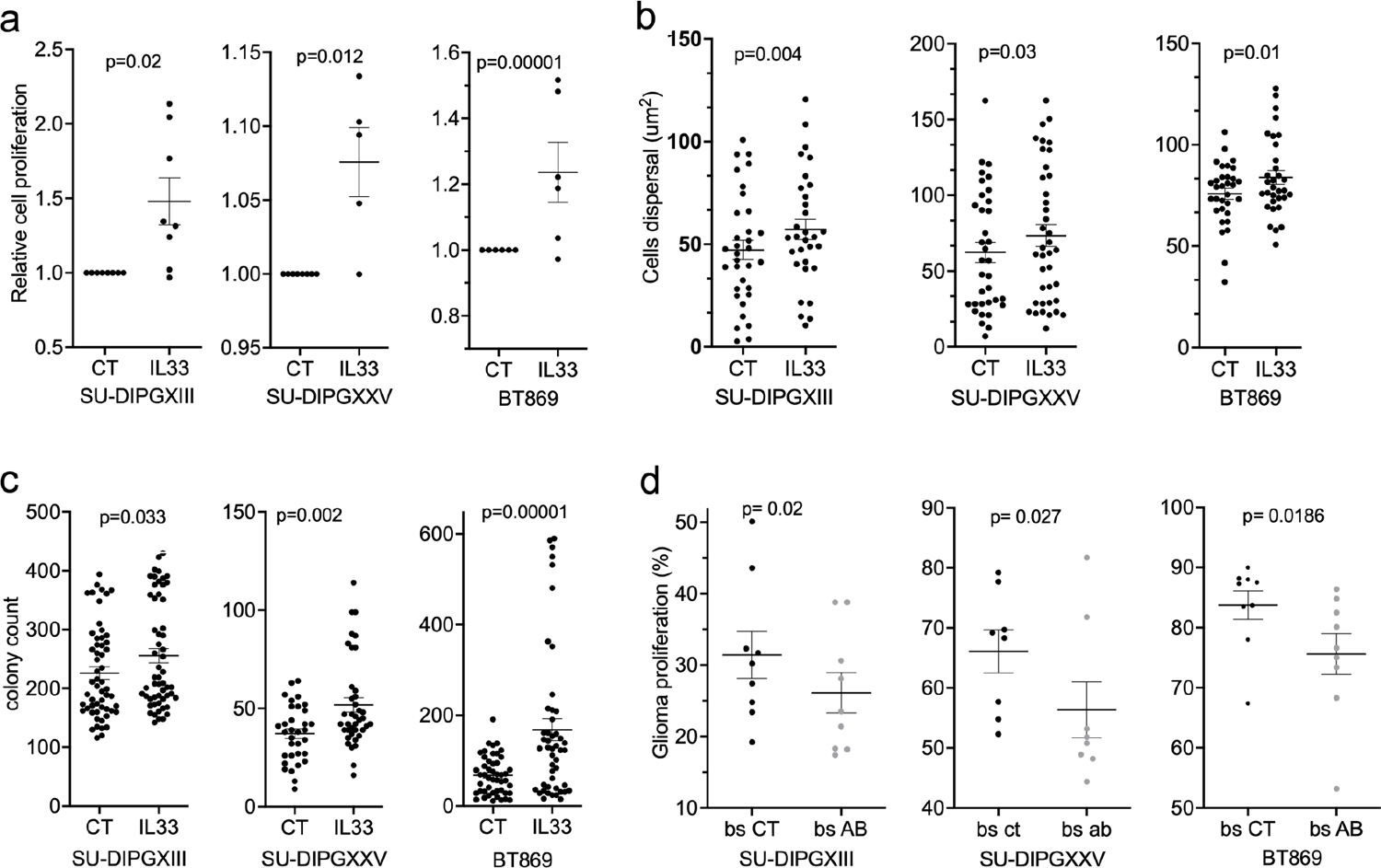
IL33 is able to impact DIPG tumorigenic properties. (a) Proliferation of DIPG cells over a period of 72hrs with IL33 (50 ng/ml) or control condition (CT). Proliferation was measured using CellTiterGlo luminescence. The data is presented as the fold change in proliferation with IL33 treatment compared to control, each point represents the average of 8 repeats. (Wilcoxon’s signed rank test; SU-DIPGXIII n=8, SU-DIPGXXV n=6, BT869 n=8) (b) Cells’ dispersal diameter refers to the expansion of each DIPG line in a Matrigel-TSM mixture (1:1) with IL33 treatment, in comparison to the CT. The data was obtained by placing a concentrated droplet of DIPG cells in the center of each gel-TSM droplet and measuring the area of the DIPG droplet after 2hrs (for a time zero measurement), and then comparing to the final area after 7 days. Images were obtained and the area measured, using ImageJ. (Fisher’s Combined Probability Test; SU-DIPGXIII n=6, SU-DIPGXXV n=6, BT869 n=5. Within each trial, a Mann Whitney U test was performed to obtain the P-values, n=7 droplets. One tailed). (c) DIPG colony count per well, after a treatment period of 5-7 days with IL33 in comparison to control. Quantification was performed with Incucyte® S3 Live-Cell Analysis system. (Fisher’s Combined Probability Test; SU-DIPGXIII n=11, SU-DIPGXXV n=8, BT869 n=6. Within each trial, a Mann Whitney U test was performed to obtain the P-values, n=6-8 wells. One tailed). (d) mIL33 neutralization trial in ex-vivo slice cultures. FACS analysis of CellTrace dye staining of DIPG cells, injected with 1μg/ml of mIL33 antibody or isotype control antibody. The cells were injected into two sides of the brainstem from a single P2 mouse brain, then slice cultures were prepared. After 72h, the proliferation of DIPGs was measured. (Wilcoxon’s signed rank test, one tailed; SU-DIPGXIII n=10, SU-DIPGXXV n=10, BT869 n=11, when n represents a single brain).

To test the role of IL33 in proliferation in the brainstem, we used CellTrace™ labeled DIPG cells injected into slice cultures, together with either IL33 blocking antibody or isotype control antibody (Fig 4d). All three DIPGs tested proliferated significantly less in the presence of the IL33 antibody (Fig 4d). The relatively modest effect might indicate limited accessibility of the antibody or perhaps the contribution of other factors present in the brainstem.

### Expression of IL33 receptors in DIPGs

We explored the *PedcBioPortal* and found that expression levels of IL1RL1, IL33’s binding receptor, are higher in DIPG and cortical HGG biopsies compared to expression levels in cortical low grade cortical gliomas (LGG) (Fig5a), a result consistent with other studies in adult HGGs in which IL33 has been suggested to play a role^26–28,31,32^. Consistent with the biopsy data, IL1RL1 was expressed at much higher levels in SU-DIPGXIII cells injected into the brainstem (organotypic slice culture) as compared to cultured cells, while IL1-RAcP (IL33’s co-receptor) was also affected, but more moderately (Fig5b). Furthermore, both receptors were expressed at higher levels in cells injected into organotypic slice cultures of the brainstem in comparison to the cortex (Fig 5c, d), and on brainstem microglia (Fig e). To test the role of IL1RL1 signaling in DIPG proliferation, a knockdown approach was utilized. SU-DIPGXIII was treated with shRNAs targeting IL1RL1, and the effect was validated by qPCR (Fig. S12). The knocked down cells were then tested in an organotypic proliferation assay (Fig 5f). Consistent with the IL33 blocking experiments, targeting IL33’s receptor reduced proliferation in the brainstem while, as expected, had no effect on proliferation in the cortex.

**Figure 5:**
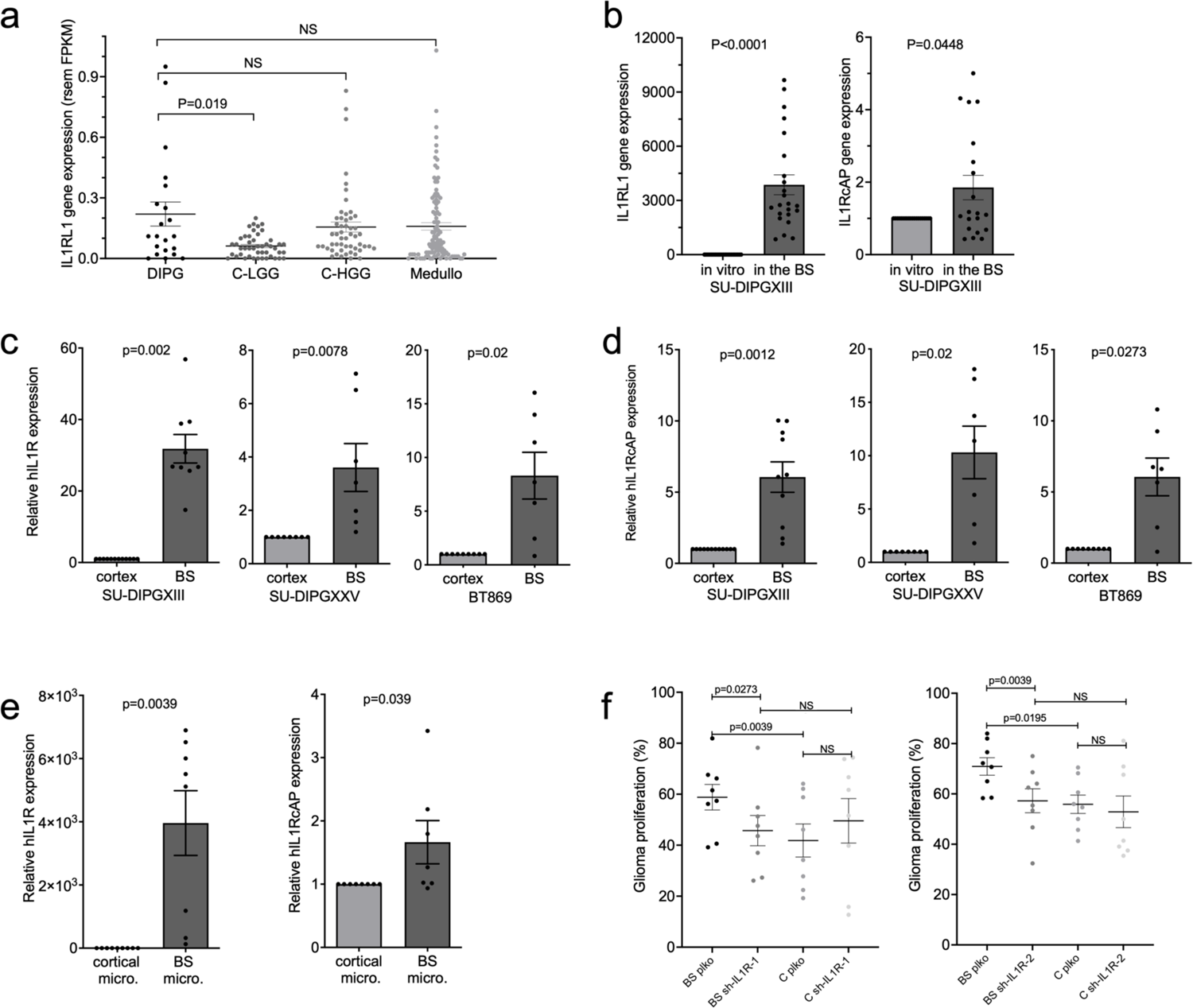
IL33’s receptors are involved in glioma-brainstem crosstalk. (a) Human IL1RL1 gene expression data obtained from PedcBioPortal. The gene’s expression within biopsies from DIPG samples was compared to cortical low and high grade gliomas and to Medulloblastoma biopsies. Each point on the graph represents a single human sample, FPKM= fragments per kilobase of exon per million mapped fragments, RSEM= RNASeq by Expectation-Maximization (Mann Whitney U test was performed). (b) qPCR analysis of IL1RL1 and IL1RAcP expression in SU-DIPGXIII cells, cultured for 72h or injected into mouse brainstems and incubated for 72h. (Wilcoxon’s signed rank test, one tailed; IL1RL1n=24 and IL1RcAP n=20). (c+d) qPCR analysis of IL1RL1 (c) and IL1RcAP (d) expression in glioma cells which were injected into the brainstem or cortex of mice, and then incubated in slice culture for 72h (Wilcoxon’s signed rank test, one tailed; SU-DIPGXIII n=9, SU-DIPGXXV n=8, BT869 n=9). (e) qPCR analysis of IL1RL1 and IL1RcAP expression in SU-DIPGXIII cells incubated for 72h on purified microglia cultures from either the brainstem or the cortex (Wilcoxon’s signed rank test, one tailed; IL1RL1 n=9 and IL1RcAP n=8). **For all qPCR analyses, the values are expressed as the average fold change in the brainstems in comparison to the corresponding cortexes ± SEM. (f) IL1RL1 knockdown’s effect on SU-<u>DIPGXIII proliferation in ex vivo slice culture</u>. FACS analysis of CellTrace dye staining of GFP+ SU-DIPGXIII cells which underwent IL1RL1 knockdown using two separate shRNAs, 72h post injection into the cortex and brainstem of the same brain. For each brain, a parallel injection of control (plko) shRNA treated cells was performed on the other side. Each point on the graph represents the percentage of GFP+ cells which underwent division (at least one), out of the entire GFP+ cell population ±SEM. (Wilcoxon’s signed rank test, one tailed; sh-IL1R-1, sh-IL1R-2 n=8).

## Discussion

In this study, we sought to begin exploring the mechanism underlying the observed spatiotemporal pattern of pediatric HGGs. Two explanations have been put forward in the past. One explanation suggests that the locations of pHGGs coincide with waves of myelination in the childhood and adolescent brains and so reflects tumor intrinsic factors, while a second possibility suggests that this pattern is due to the microenvironment. These two explanations are not necessarily mutually exclusive.

Here we tested the potential of the brainstem surroundings to differentially impact DIPGs’ survival and proliferation. As compared to the cortex, the brainstem shows a potential to better support the progression of HGGs of brainstem origin. This was demonstrated by the increased survival and proliferation of DIPG tumors in the brainstem. Our findings supports the idea that at a young age, the microenvironment of the brainstem is better able to foster the progression of DIPG tumor cells. Interestingly, these differential effects were in most cases specific to DIPGs as compared to pediatric cortical HGG lines, suggesting a cell-intrinsic component of the tumor cells.

The pro-tumor activity of microglia and macrophages is well documented in adult gliomas^23^. As in other gliomas, microglia/macrophages account for a significant portion of the cellular mass of DIPGs^33^. Although the involvement of microglia in promoting DIPG propagation may not be surprising in itself, the differential ability of brainstem microglia as compared to cortical microglia to promote DIPG growth, is striking. In recent years there has been accumulating evidence indicating differences in microglial populations in different anatomical locations, including RNA transcription^6,34^ and functional differences ^35–37^. Here we show that microglia differentially support cancer cells based on their anatomical location, partly through IL33.

Accumulating evidence suggests a role for IL33 in adult glioma. Higher expression of IL33 in human glioma specimens have been reported ^26,27^ and is associated with poor prognosis^28,32^. IL33 has been shown to act directly on glioma cell migration, invasion, and growth^26–28,31^. The role of IL33 in pediatric HGGs has not previously been explored. Based on our findings here and on published data, IL33 is more highly expressed in the brainstem and particularly in the pons of young mice as compared to the cortex (from P2 to P23) ^38^. According to the Allen Brain Atlas, by day 56 there are no longer any spatial differences in IL33 expression in mice ^39^. Also, IL33 is more highly expressed in childhood DIPG biopsies, possibly indicating similar expression patterns in humans and a tumorigenic role. Furthermore, IL1RL1’s expression in DIPGs is upregulated in brainstem slices, and specifically by brainstem microglia, while inhibiting IL33 and IL1RL1 decreases pro-tumorigenic effects in an anatomically specific manner. All of this suggesting a signaling system that is modulated by the native tumor surroundings. Together, the expression data and the impact on DIPGs in culture and in brainstem slices, makes IL33 a likely factor in the brainstem’s micro-environment that contributes to the affinity of DIPGs to this location. Future testing of IL33 as a target using *in vivo* mouse models will be needed before considering reagents such as IL33 blocking antibody ANB020 as a possible therapy in DIPG cases.

## Acknowledgments

We thank the Koch Institute’s Robert A. Swanson (1969) Biotechnology Center for technical support, specifically the Bioinformatics Facility. We also thank the Hebrew University, Faculty of Medicine Core Research Facility and particularly to Zakhariya Manevitch for valuable help with the two photon microscopy. We thank Keith L Ligon (Dana-Farber Cancer Institute) for BT869, and Michelle Monje (Stanford University School of Medicine) for the SU-pcGBM2, SU-DIPGXXV, and SU-DIPGXIII. We are grateful to Dr. Norman Grover (Department of Experimental Medicine, The Hebrew University) for helpful advice regarding the statistical analyses.

