## Supplementary methods and figures for "An affinity for brainstem microglia in pediatric high grade gliomas of brainstem origin"

**Animals**

C57BL/6 mice were obtained from ENVIGO (Rehovot, Israel). Animal handling adhered strictly to national and institutional guidelines for animal research and was approved by the Ethics Committee of the Hebrew University. Mice both males and females between day 2 and 4 after birth were used.

**Antibodies**

Anti-mouse IL33 AF3626 (15ug/ml) and human IL33 AF3625 (15ug/ml) (R & D systems, Mi, USA), Iba1 (CP290B, Biocare Medical, Pacheco, CA, USA, 1:500), actin (ab8224, Abcam, Cambridge, UK,1:1000), CD11b (deposited to the DSHB by Springer, T.A.; DSHB Hybridoma Product M1/70.15.11.5.2,1:20), Anti-human nuclei clone 235-1 (Merck, Millipore, Burlington, MA, USA,1:60), TMEM 119 (ab209064 Abcam, Cambridge, UK,1:800).

**Patient-derived tumor cell cultures**

All cultures were maintained as neurospheres in tumor stem medium consisting of Neurobasal(-A) (Invitrogen), B27(-A) (Invitrogen), human basic fibroblast growth factor (bFGF) (20 ng/ml) (Reprokine), human epidermal growth factor (EGF) (20 ng/ml) (Reprokine), human platelet-derived growth factor (PDGF)–AB (20 ng/ml) (Reprokine), and heparin (10 ng/ml).All cell culture models were received from Michelle Monje(1,2) (Stanford University School of Medicine), and validated by short tandem repeat (STR) DNA fingerprinting and tested for mycoplasma. Information about age, sex, and clinical characteristics associated with each cell culture can be found in table below:

| Patient derived cell lines | Tumor type, location and grade | Age at diagnosis (years) | Sex | Histone – mutational status | Prior therapy |
|---|---|---|---|---|---|
| BT869 | Anaplastic Astrocytoma NOS, Midline Brain stem | 7 | F | H3.3K27M | none |
| SU-DIPGXXV | DIPG, pons, WHO grade IV | 4 | F | H3.3K27M | XRT |

| | | | | | |
|---|---|---|---|---|---|
| SU-DIPGXIII | DIPG, pons, WHO grade IV | 6 | F | H3.3K27M | XRT |
| SU-pcGBM2 | Pediatric cortical gliobastoma, WHO grade IV | 15 | M | wt | none |
| KNS-42 | Pediatric cortical gliobastoma, | 16 | M | wt | |

***Preparation of glia primary cell cultures:***

Brainstems and cortices of 2-3 day old mice were used. A minimum of 3 cortices or brainstems were combined per dissection. After dissociation, the cells were treated to produce either astrocyte or microglial enriched cell cultures.

***Astrocyte primary cell culture:***

Briefly, the main modifications are the addition of EGF (100ng/ml) in the first two days of culture and an AraC (0.1mM, Sigma Aldrich) selection cycle once the cells achieve confluence. Representative cultures were stained with anti-S100b to test astrocyte content (at least 95% purity was achieved).

***Microglial-enriched primary cell culture:***

High density glial cultures obtained from cortex and brainstem were grown with gmCSF **(**0.5ng/ml**).** Once the cultures became confluent, circular cells began to float in the medium, which are microglia. At this point cultures were treated with 2.4 mM lidocaine for around 15 minutes, which enhanced the microglia's ability to de-attach from the rest of the culture. The cells were then plated in a density of about 20,000 cells/$cm^2$.Cultures were stained with anti-Iba1 to test for microglial content (approximately 85% purity was achieved).

**Mixed glia cultures:**

Were prepared similarly to microglia, but with no factor treatment. These became confluent within one week, and ready for trial.

**Freshly isolated cortical and brainstem cells:**

Similarly to the initial stages of microglia and astrocyte preparation, cells were dissociated from brain tissue using trypsin B, followed by gentle trituration. The cell suspension was then plated on PDL coated slides and fixed withing two hours of dissociation. This rapid fixation is needed to preserve the in-vivo microglial- specific gene expression. After fixation, cells were stained with TMEM119, CD11b, or mIL33, as described in the 'immunostaining' section.

**GFP+ or luciferase DIPG cell line preparation:**

eGFP (Fuw originated-puro-2A-EGFP) or Luciferase (LentipLVX-Luc-Puro) plasmids were packaged together with helper plasmids (pΔ8.9 and PDMG) to generate replication-deficient lentiviruses from adherent 293T cells. Each human patient derived primary high-grade glioma cell line was infected with Lenti-GFP/Luciferase viral supernatant and allowed to recover for 1 week. GFP-positive cells were isolated for purity by FACS (BD FACS Aria) and returned to culture. Luciferase positive cells were isolated via puromycin (1ug/ml) selection. Validation of the sorting was done using FACS analysis and by observing the cell fluorescence under the microscope (with at least a 90% positive GFP rate).

**Ex-Vivo Injections:**

Tumor cells (10,000 cells/µl) were re-suspended in Matrigel (Corning, NY, USA) and TSM (1:1) and injected (1µl) using Narishige MM-3 micromanipulator (Tokyo, Japan) into the cortex and brainstem of mice, two injections per region. The injections of each region were performed consecutively, alternating the order of injections between brainstem and cortex. After injections, the brain regions were gently separated and were cultured as described before (3) and in the next section

**Slice Cultures:** The tissue pieces were incubated on top of 40µm cell filters. Each well was filled with medium (Neurobasal, 15mg/ml glucose, AmphoB 25 µg/ml, Pen-Strep, and B27 (without vitA)) up to the level of the filter, as to allow diffusion. Incubation was for 24-72h in 5% $CO_2$, $37^0$C.

***Ex- vivo Multiphoton imaging and quantification***

Each injected cortex and brainstem pair was incubated for 24hrs, then sealed with a cover slip (moisturized with PBS 1X) and imaged, using the large image option of *Nis-Elements*, and the maximal width in the Z stack option so as to capture all of the injected GFP+ cells. The GFP+ cells were quantified in *Nis-Elements* using the 'sum measured area' ($\mu m^2$) of all GFP positive areas, of each image (representing the sum of the cell amount per z slice, per image). To quantify the dispersion of cells in each slice, the maximal area (length X width) of GFP+ cell distribution was manually measured using *Nis-Elements* tools, and this area was multiplied by the maximal Z stack height in which cells were found ($\mu m^3$).

**DNA quantification using qPCR:**

Testing cell number with DNA based qPCR and primers was carried out based on a previous paper (4) with some modifications. Briefly, DNA from a controlled number of glioma and mouse (astrocytes and microglia) cells was acquired, then a qPCR reaction (iTaq Universal SYBR Green Supermix, BioRad, Rishon Le Zion, Israel) using either a human specific primer or a primer designed to detect both human and mouse cells, was performed on the control DNA samples. The results were used to create transformation formulas from cycle threshold (CT) to cell number/DNA amount with a high prediction power ($R^2$=0.99 for both primers). To test the strength of these transformation formulas, sets of quantified human and mouse DNA samples (from 100% human DNA, 90% human with 10% mouse, etc., all the way to 100% mouse DNA) were tested. Using our transformation formula, we were able to accurately estimate the ratio of mouse and human cells in a mix culture, with a strong predictive value ($R^2$=0.98).

**Co-culture proliferation experiments:**

8,000 human HGGs were placed on top of mouse glial cell cultures (4$cm^2$ wells)- astrocytes and microglia, originating either in the brainstem or the cortex of the same mice. For each condition, within each trial, T0 wells were acquired about 2 hours after the trial began. The co-culture was then incubated for 72 hours in glioma medium, after which DNA was acquired using the Monarch DNA cleanup kit (NEB, Ipswich, MA, USA), and DNA-based qPCR quantification was done, as described above. The analysis was performed by normalizing each sample's glioma cell quantification (human primer read) to the total cell quantification (mouse and human cells), to produce the ratio/proportion of glioma cells within the co-culture. Then the fold change in the glioma's ratio within the culture at time=0 in comparison to the ratio recorded under the same conditions 72 hours later was calculated to evaluate the effect.

**luciferase based viability assay**

Luciferase labeled DIPG cells were plated on top of purified microglia or astrocytes (670 cells/well), in 0.3$cm^2$ wells. Following a 72h incubation, luciferase activity was measured using LUMIstar® Omega (BMG LABTECH) microplate luminescence reader. Each experiment consisted of a minimum of four wells per condition, and each experiment was repeated at least six time per cell line.

***Microglial conditioned medium's effect on DIPGs' clonability:***

For each trial a confluent microglial cell culture was prepared, taking care to maintain similar conditions for the brainstem and cortical derived microglia. Once cell cultures reached similar confluence, the cells were thoroughly washed to remove any residual serum, and TSM without factors was added. For each 12well well, 1.5ml of medium was added. TSM was incubated with the microglia for 72hrs, after which the medium was collected, filtered, 12.5% factors and 100% B27 were added, and the three DIPG cell lines were incubated with the prepared conditioned medium in the following manner. 400 cells of each of the DIPG cell lines were plated in 0.3cm$^2$ wells, with about 80ul conditioned medium (CM). This low density plating allowed clear segregation of colonies. For each condition BS CM and cortical CM, at least 7 wells were plated. After 5 days incubation the cultures were analyzed using Incucyte® S3 Live-Cell Analysis system (whole well image X4). This data was then statistically analyzed.

***Preparation of samples for RNA-sequencing:***

Brainstem and cortical microglia were each cultured with DIPGXIII for three days before RNA was extracted. Each sample was a separate independent trial (dissection/cell culture). In order to preserve RNA integrity, a trizol-based RNAKIT-Direct-zol™ RNA MiniPrep (Zymo Research) kit was used and the acquired RNA was placed in liquid hydrogen or at -80°C degrees at all times. All samples had a 260/280 OD of at least 1.93 (and an average of 2), and a RIN value of at least 9.3.

**RNA Data processing:** We utilized STAR, RSEM, Tximport, and DESeq2 to analyze the FASTQ raw RNA-seq data. We used STAR (version 2.5.3a) to align the reads to the Hg38 human and Mm10 mouse reference genomes (5). We then used RSEM (version 1.3.0) to generate BAM files containing estimates of the gene and isoform expression levels from the single-end reads (6). We conducted hierarchical clustering with Euclidean distance calculations (MATLAB 2018b) using log2 (FPKM) values.

**Data analysis:** We utilized the R package Tximport (version 1.8.0) to generate a count matrix from the RSEM output (7). DESeq2 does not accept reads of length 0; we assigned a length of 1 to these entries and filtered out any genes with no reads across the samples in order to optimize the process. We then used this modified count matrix for DESeq2 analysis (version 1.20.0) (8). To identify differentially expressed genes (DEGs), we identified pairs of experimental groups for comparison. We set the false discovery rate (i.e. alpha) to 0.05 and used Benjamini-Hochberg FDR-controlling correction method in this analysis. DEGs were defined as those with abs(log2FoldChange)>1 and padj<0.05.

To obtain the subcellular localization information we used the UniProt Knowledgebase (UniProtKB) (9). Ensembl identifier and gene names were used as search input. Only annotations from reviewed entries (UniProtKB/Swiss-Prot) were included, and the two lists, resulting from Ensembl ID search and gene name search respectively, were combined following annotation.

**mIL33 expression quantification:** Immuno-staining trials (tissue and microglia culture) were performed in Nis-elements, using the ratio of mIL33 positive 'sum area' within the CD11b/Iba1 positive 'sum area' analysis (representing IL33's expression with microglia cells).

**Western blot analysis-** protein was extracted from the various DIPG cells/tissue samples using a lysis buffer: 4% SDS, 50mM Tris, pH=7.4. The samples were separated on a 10% gel and transferred to a PVDF membrane (Immobilon P, Millipore, Burlington, MA, USA) for tissue samples, or to a nitrocellulose membrane for cells. The membranes where then incubated with the primary antibodies- against mouse IL33 (R&D systems, AF3626), and actin (ab8224, Abcam, Cambridge, UK). Then the appropriate secondary antibody was used- HRP conjugated donkey anti-goat, the band imaging was performed and the statistical analysis was done using GelQuant Express analysis.

***qPCR:***

RNA was extracted using Direct-zol$^{TM}$ RNA MiniPrep (ZymoResearch) and cDNA was prepared using qScript cDNA Synthesis Kit (Quantabio). RT-PCR was performed with triplicates per sample, using Universal SYBR Green Supermix (BioRad, Hercules, CA). Differential expression was determined using the delta CT method. Each primer set was tested on human and mouse samples to make sure each primer is species specific. In each experiment an average of two control genes was used. Primers used are as follows:

qPCR primers:

| Gene | specie | Rev/Fw | Sequence |
|---|---|---|---|
| NKX6 | mouse | forward | CCGAGTCCTGCTTCTTCTTG |
| NKX6 | mouse | reverse | CTTGGCAGGACCAGAGAGAG |
| HOXB8 | mouse | forward | CACAGCTCTTTCCCTGGATG |
| HOXB8 | mouse | reverse | CAGGGTCTGGTAGCGACTGT |
| HOXB6 | mouse | forward | CAGGGTCTGGTAGCGTGTGT |
| HOXB6 | mouse | reverse | CTTCTCCAGCCGCCTCCT |
| FOXA2 | mouse | forward | GGTGAGACTGCTCCCTTGAG |

| | | | |
|---|---|---|---|
| FOXA2 | mouse | reverse | GGGAAATGAGAGGCTGAGTG |
| PHOX2B | mouse | forward | GCTCCTGCTTGCGAAACTTA |
| PHOX2B | mouse | reverse | TGAGACGCACTACCCTGACA |
| SNRPD3 | human | forward | CCTTCGCCGTAGCATCTTT |
| SNRPD3 | human | reverse | TCGGCCTCATGCAGTACTTT |
| TNC | human | forward | CAGACACAGCCACATCCTTC |
| TNC | human | reverse | GAACCTGGTGTCTTCCCTGA |
| IL1RL1 | human | forward | CCGCATCAACACAAACTTGC |
| IL1RL1 | human | reverse | CAAATTCAGGGCCAGACAGT |
| CXCR1/2 | human | forward | CGCTCCGTCACTGATGTCTA |
| CXCR1/2 | human | reverse | AAATCCAGCCATTCACCTTG |
| EMC7 | mouse | forward | AGAGCATGTCGGCTTCCTTA |
| EMC7 | mouse | reverse | GACTCGGACAGGGTCAAACT |
| EMC7 | human | forward | AGAGCACGTCGGTTTCCTTA |
| EMC7 | human | reverse | TCGAACGGGATCAAATCTGT |
| IL1RAP | human | forward | TGCATCTTTGACCGAGACAG |
| IL1RAP | human | reverse | GTTGGGGCTTAGAACAACCA |
| CHMP2A | mouse | forward | AAGGCCAGATGGATGCTGT |
| CHMP2A | mouse | reverse | CCGCATCAACACAAACTTGC |

***Recombinant IL33 concentration calibration:***

According to previous studies the optimal concentration for recombinant human IL-33 (PeproTech #200-33 10) was tested between 25-100ng/µl. We therefore decided to create a dose response curve for the effect of IL33 on SU-DIPGXIII proliferation using a couple concentrations under and over the previously reported amount (via X2 serial dilutions). For this, about 1,000 DIPG cells were plated in $0.3cm^2$ wells, in about 80ul TSM based medium+IL33 or control. 5 wells were plated per condition, and after 72 hours, the number of cells in each well was quantified by adding CellTiter-Glo reagent and quantifying the culture's luminescence.

***Recombinant IL33's effect on tumorigenesis-***

***Three dimensional growth experiments:***

The recombinant human IL33 was added to a TSM and Matrigel mixture (1:1 ratio) in a 50ng/ml concentration, with DIPG cells highly concentrated to create very dense cell-matrigel droplets. These marked a specific starting point. Control droplets were also added. After initial solidifying of the gel, the radius of the droplet was expanded by adding more Matrigel-TSM mix. After final solidification, TSM based medium was added on top of the Matrigel drops to cover them. The diameter of each DIPG drop was measured after 6 hours and compared to the final diameter after 7 days (pictures were taken and the diameter measured using ImageJ). In each trial at least 7 droplets for each condition were plated.

***Proliferation experiments:***

CellTiter-Glo reagent was used as described above in 'Recombinant IL33 concentration calibration' and more DIPG lines were added.

***Colony count and average colony size experiments:***

Trials were performed as described above in the '*Microglial conditioned medium's effect on DIPGs' clonability*' section, but this time the cells were plated with TSM based medium with 12.5% of factors+100% B27 (without vitA), with the addition recombinant IL33 (50ng/ml) or a CT condition.

***Immunostaining:***

Fixation was done with 4% paraformaldehyde for 15 min at room temperature. Three PBS washes for 5 min each were performed. Blocking solution was added for 1 hr at RT, and then diluted primary antibody was applied (in blocking solution) on each slide, and incubated overnight at 4°C. The next day, 3 PBS washes were performed, and a 1hr incubation with the secondary antibody was completed. Finally, the slides were washed again, dried, covered with Dako-mounting medium with DAPI and sealed with a cover slip.

Blocking solution: 5% goat serum (or FBS for Iba1), 0.05% Triton X-100 for tissue or 0.01% for cells, 0.001% $NaN_3$ in PBS.

**mIL33 neutralization- control condition**

An additional cortical injection was used to exclude any brains in which no brainstem preference was detected from the statistical analysis, excluded brains: DIPGXIII 3/13, DIPGXXV 1/12, BT8693/14.

***IL33's receptors gene expression in slice culture and in co-culture:***

For DIPG IL33's receptor expression (IL1RL1 and IL1RAcP), DIPG cell lines were injected into the cortex and brainstem of 2-3 day old mice and slice cultures were kept for 72 hrs. At least 7 brains from each cell line were injected. After 72 hrs, RNA was extracted from each brainstem-cortex pair, and qPCR was performed using the appropriate human primers.

For the receptor expression in co-culture, DIPG cell lines were incubated with mouse microglia from the BS and cortex for three days. After this, RNA was acquired using the same kit and qPCR performed for both human and mouse genes.

**IL1RL1 knockdown:**

For IL1RL1 knockdown experiments we used Mission® shRNA PLKO lentiviral vectors which include puromycin selection marker: TRCN0000358832/sh-IL1R-1, and TRCN0000058513/sh-IL1R-2. For a control we used an empty vector (pLKO.1-puro), all vectors purchased from Sigma Aldrich. Lentiviral infections: Prior to infection each virus was concentrated with a 4 hour, 4 degree, maximal speed centrifugation, in 4X buffer (EDTA 0.5mM, NaCl 100mM, Tris 50mMM Ph 7.4, sucrose 100mg/ml in DDW). SU-DIPGXIII cells were infected with each of the three concentrated viruses. After a couple of days (once the cells appeared stable), puromycin selection was applied, 1µg/ml, and after three days an increased dosage was applied, 2µg/ml, for a total of a week. Each shRNA infected SU-DIPGXIII cell line was then tested for the degree of knockdown using qPCR.

**PedcBioPortal** Gene expression data analysis

The expression level of the genes IL33 and IL1RL1 in DIPG patients were compared with their expression level in low grade glioma patients, high grade glioma patients and medulloblastoma patients. The patient samples included in every comparison group were defined based on the clinical information included in CANCER_TYPE, CANCER_TYPE_DETAILS and TUMOR_TISSUE_SITE columns. Every sample group was built by intersection of characteristic values included in these columns. Details of the values used as selection criteria to build the 4 comparison groups can be found in the supplementary PedcBioPortal file group_criteria.xlsx. For patients that had duplicate samples, the first sample was selected based on sample number (i.e., the sample with lower number) and all other samples were removed. If duplicate samples included samples labeled with "CL-adh" (corresponds to the adherent FBS cell lines) and samples labelled "CL-susp"(corresponds to serum free cell lines) the samples marked with "CL-adh" were removed.

Statistical significance of the comparisons was determined using the one-sided Mann-Whitney U test. The scripts for the analysis were written in python 3.7[3] using SciPy package[4].

**Statistical Analysis:**

The design of the experiments throughout the paper generally compared two conditions, providing an intrinsic control within the experiment and making the comparisons coupled to one another. In all the experiments, averages of intrinsic repeats were compared. In addition, as the amount of repeats per trial (n) was about 6-12, the population compared was nonparametric. Henceforth, the appropriate statistical test was a one-tailed Wilcoxon signed rank test. There were a few exceptions. First, in the 3D growth assay a Mann-Whitney U Test was used to obtain a p-value within each trial (which contained 7 repeats per condition) in order to compare the uncoupled droplets, and then the Fisher's combined probability test was performed to combine the p-values of all the trials. Secondly, in the colony count and average size experiments, a similar technique was applied. An intrinsic Mann-Whitney U test was performed per trial (8 repeats per condition) in order to compare the uncoupled wells of each condition, and then the Fisher's combined probability test was performed to combine the p-values from all of the trials. And lastly, for all immuno-staining analyses, for which the appropriate test is uncoupled, a Mann-Whitney test (with the Fisher's combined probability test when needed) was also executed. In addition, Bonferroni correction was performed for each trial in which multiple comparisons were performed. For the Mann-Whitney U Tests and the Wilcoxon signed rank tests, GraphPad Prism 8.4.2 version was used. The Bonferroni correction was manually added and calculated per each appropriate P-value. For the Fisher's combined probability test, the Chi-square score was manually calculated using the p-values obtained from the Mann-Whitney analysis, using the following formula: $-2\sum_{i=1}^{k}\ln(p_i)$, where $p_i$ is the p-value for the ith hypothesis test, k is the number of tests being combined and the degrees of freedom=2k. Using the calculated Chi square score and the degrees of freedom per set of trials, we used an online calculator from Social Science Statistics, to calculate the combined p-value. (https://www.socscistatistics.com/pvalues/chidistribution.aspx).

Bibliography

1. Venkatesh HS, Johung TB, Caretti V, Noll A, Tang Y, Nagaraja S, et al. Neuronal

Activity Promotes Glioma Growth through Neuroligin-3 Secretion. Cell. 2015 May 7;161(4):803–16.

### Supplementary Figures

**Supplementary Figure 1: a detailed presentation of the GFP+ cell count after 24hr incubation in slice culture.** Quantification was done with *FACS analysis. (a) For each glioma, each separate trial (different cell passage, indicated by a number on the graph) is presented with individual values, coupled according to the brain of origin. Both factors, cell passage number and a separate brain, affected inter-trial variation (b) For each glioma, the standard deviation per trial is portrayed.*

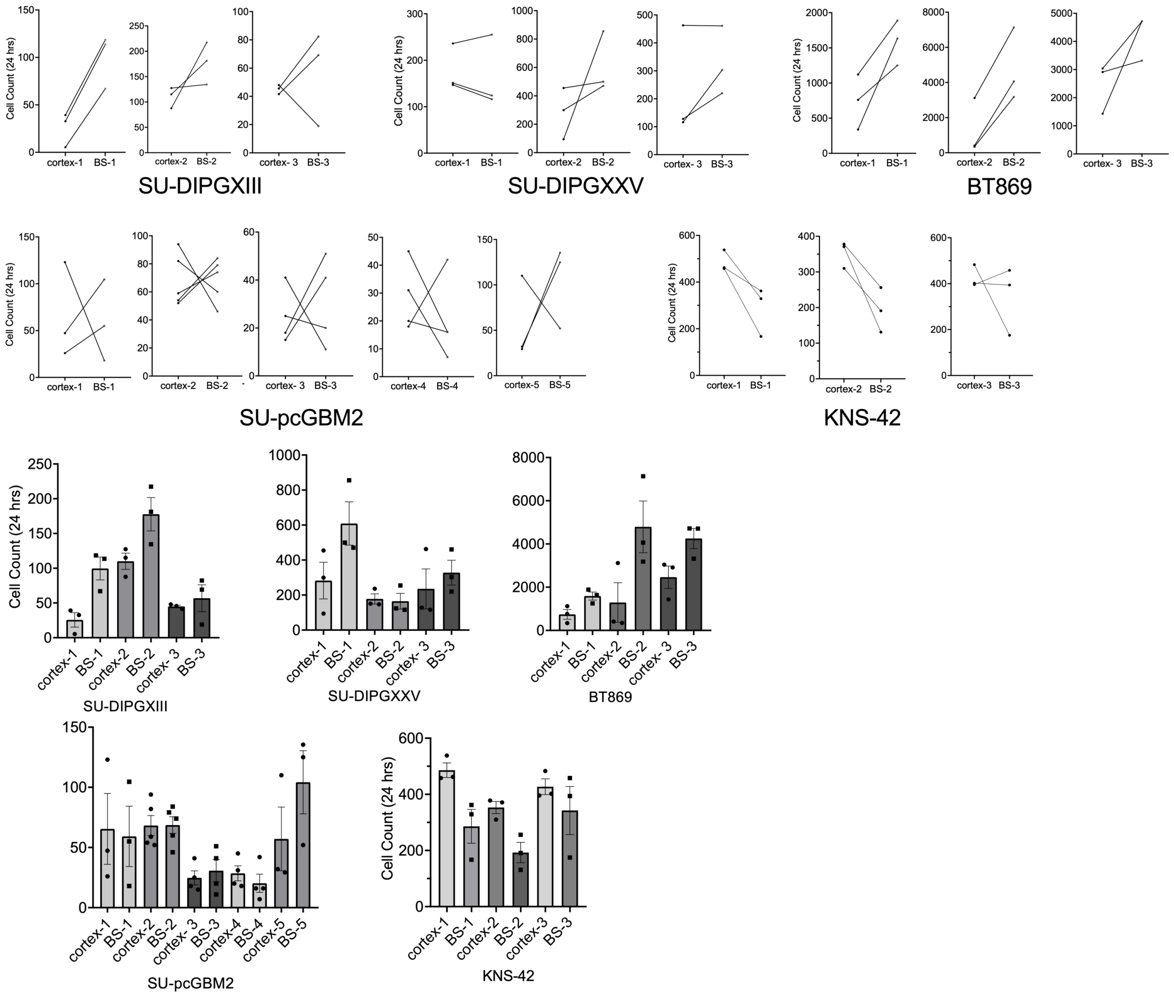


**Supplementary Figure 2:**

*FACS analysis of PI staining of GFP+ glioma cells 24 hrs post injection into the cortex and brainstem of the same brain. Each point represents the percentage of PI+ cells out of the GFP+ population, indicating glioma cell death ±SEM. (Wilcoxon's signed rank test, one tailed; -DIPGXIII n=10,BT869 n=10, SU-DIPGXXV n=10, SU-pcGBM2 n=9, KNS-42 n=9).*

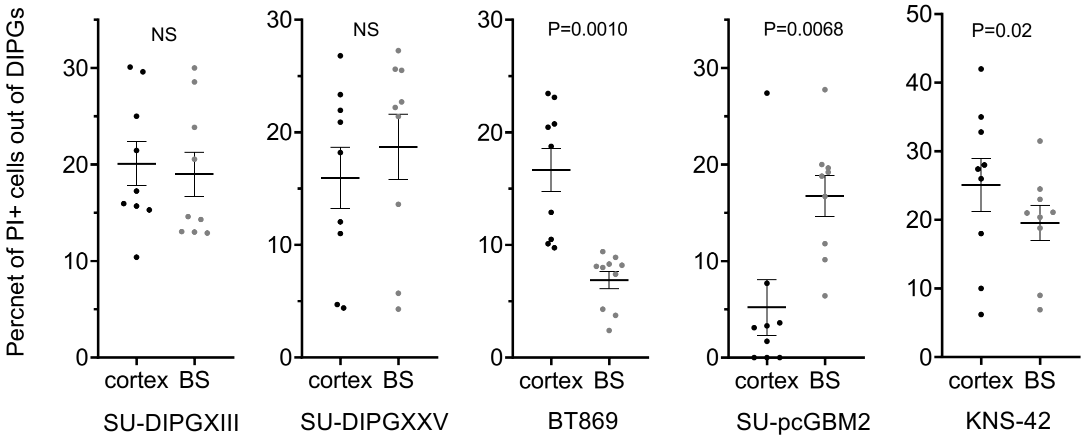


**Supplementary Figure 3:** a detailed presentation of the CellTrace proliferation quantification after 72hrs incubation.

*(a) For each glioma, each separate trial (different cell passage, indicated by a number on the graph) is presented with individual values, coupled according to the brain of origin. Both factors, cell passage number and a separate brain, affected inter-trial variation (b) For each glioma, the standard deviation per trial is portrayed.*

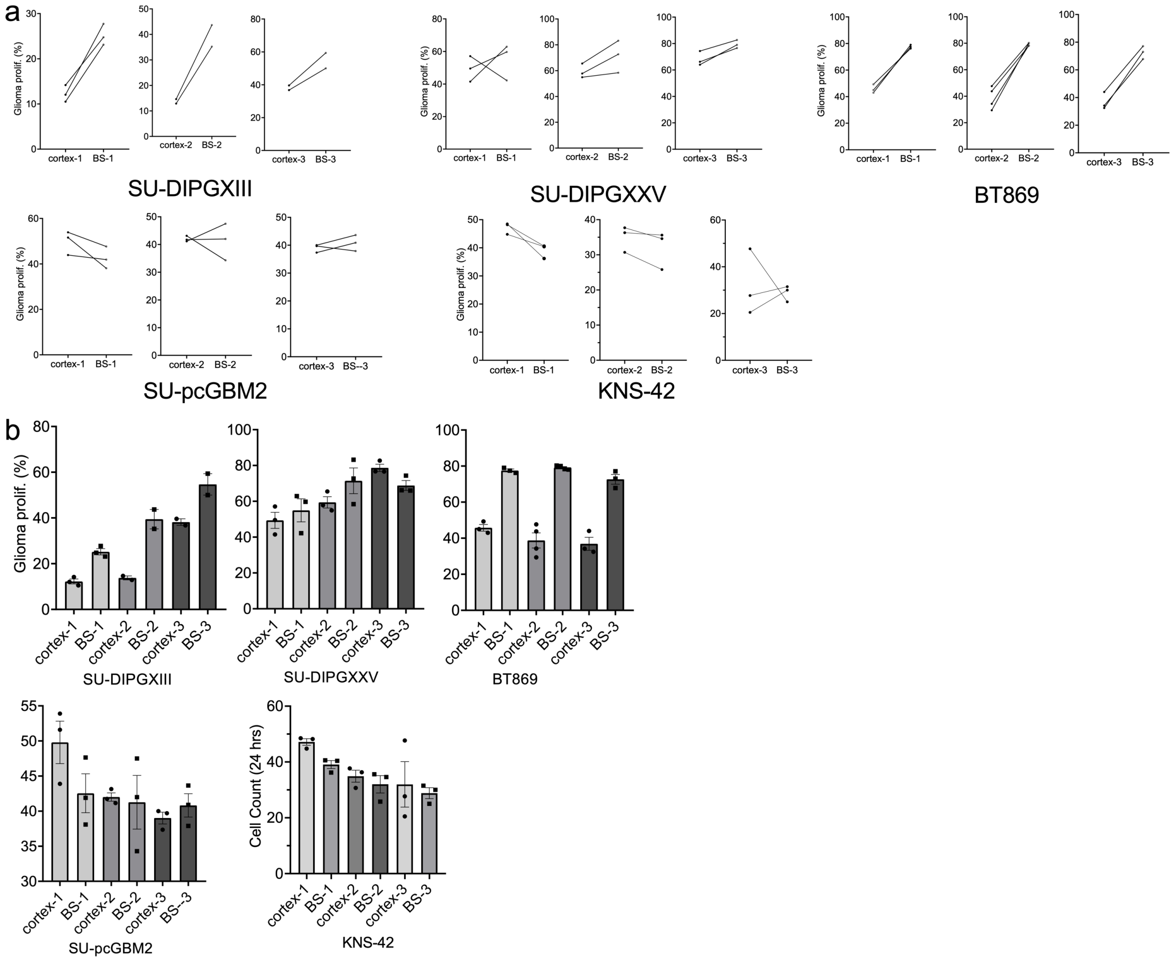


**Supplementary Figure 4: Calibration of qPCR DNA/cell number quantification:**

(a) DNA from a predetermined number of DIPGXIII cells was acquired, then a qPCR reaction using a human specific primer was performed. The results were used to create a correlation between the number of cells within the sample to the cycle threshold (CT). This produced an exponential relationship between the number of cells and the CT with a strong predictive power ($R^2$=0.99). This formula was later used for transformation from CT to cell number/DNA amount in glioma proliferation trials.

(b) In order to test the strength of this transformation formula, sets of mixed human and mouse DNA samples were created (which resembled the co-culture system) in a controlled manner (from 100% human DNA, 90% human with 10% mouse, 80% human with 20% mouse etc, all the way to 100% mouse DNA). Then the transformation formula was used to try and predict the CT outcome according to the human DNA amount within the mix. A linear relationship between the predictive values and the actual CT outcome from the qPCR reaction was observed, with a strong predictive value of $R^2$=0.98. This tested the transformation formula's strength, and reinforced the confidence in the ability to transform CT to DNA quantification.

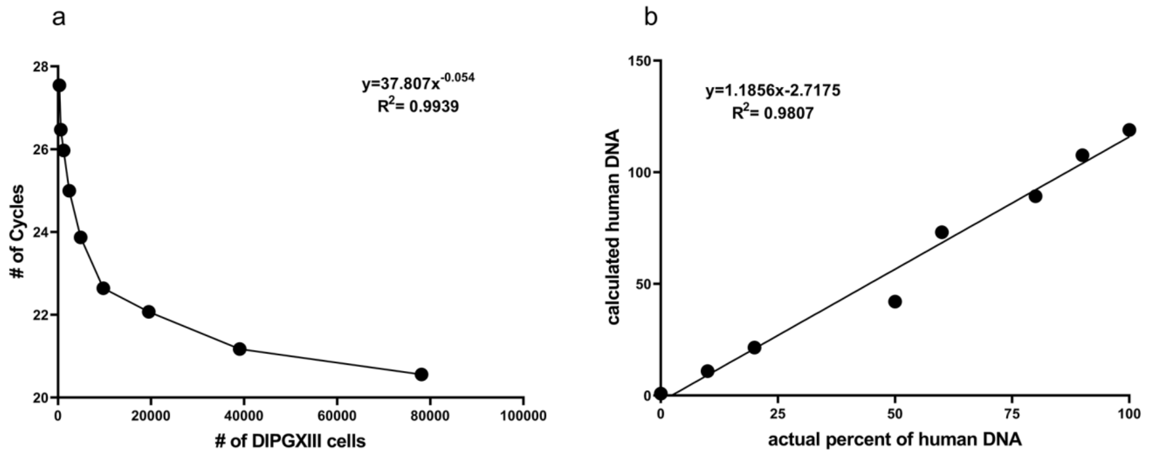


**Supplementary Figure 5:** a detailed presentation of the glia-glioma co-culture glioma proliferation trials, presented as the fold change from T0 measurement. The separate trials are individually presented with an indication of either the same origin brains (indicated by the connective BS-cortex lines), as well as marked (roughly) for the separate cell culture passage for each line (indicate by P1-4). Both factors affected inter-trial variation. (a) presents microglia co-cultures, (b) astrocytes, and (c) for the luciferase assay. Each point is the average of at least 3 repeats.

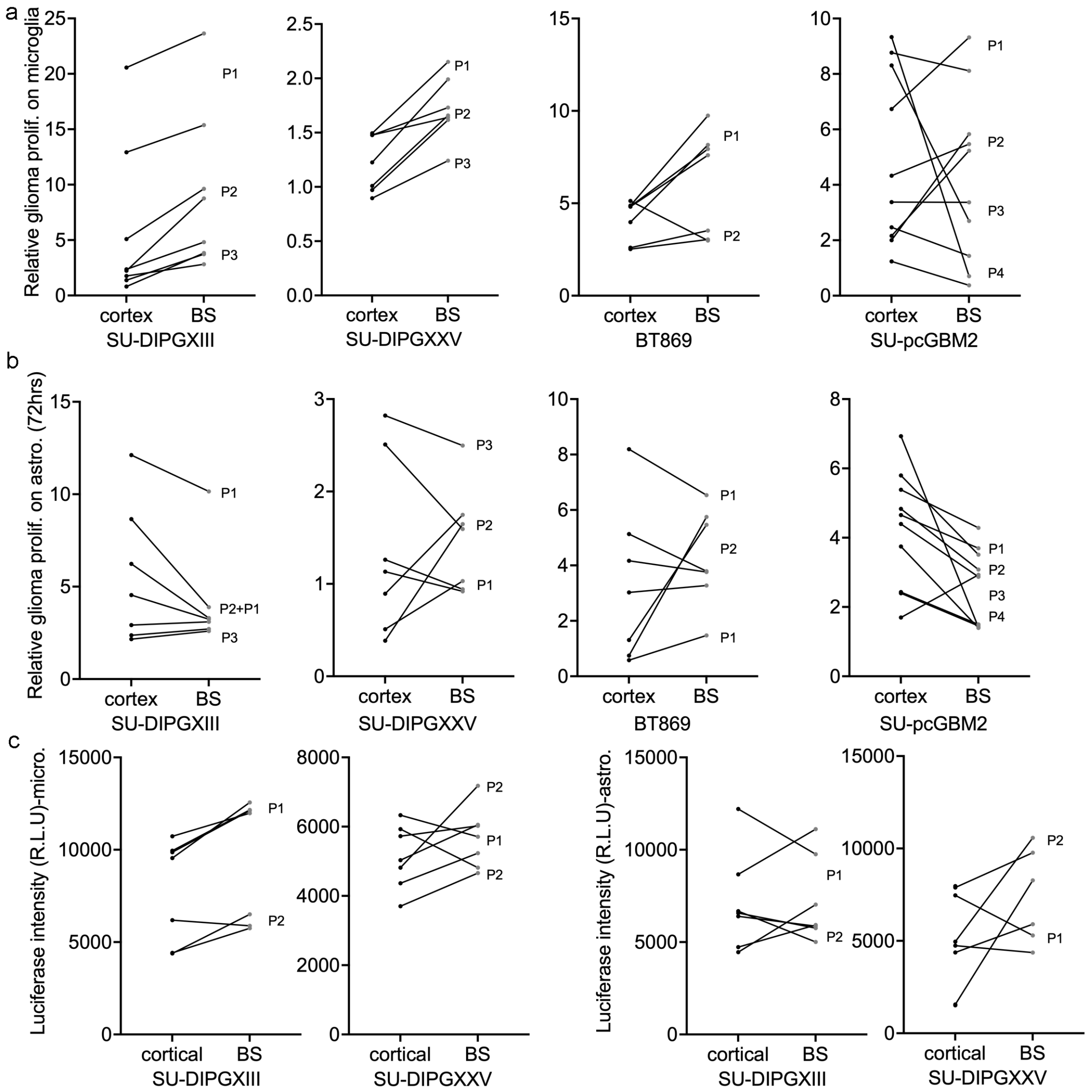


**Supplementary Figure 6: quantification of DIPGXIII grown over microglia**

DIPGXIII cells (4000) were plated over cortical and brainstem microglia slides and grown for 72h, after which fixation and staining with human anti-nuclei antibody was performed to mark the cancer cells. The number of cells on each slide was quantified manually via Nis-elements, at least 10 random fields were imaged per slide, and 6 slides were made per condition per trial. Each point on the graph represents the median of 8-10 fields imaged on the slide. (Wilcoxon's signed rank test, one tailed; n=6). The graph indicates brainstem-cortex cell cultures originating from the same brains.

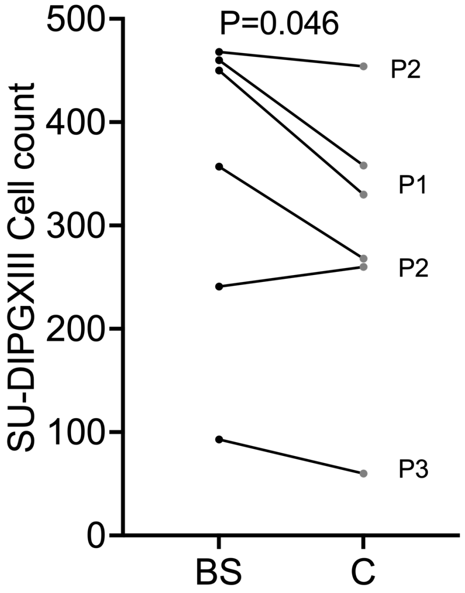


**Fig S7: expression of IL33 in freshly isolated cortical and brainstem cells**

Freshly acquired brainstem and cortical cells, were plated on slides and fixed after no more than two hours, followed by staining for specific markers.

a) Slides were stained with TMEM119 (specific microglial marker) and for CD11b (a common myeloid marker). Analysis was performed with Nis-elements by measuring the percent of TMEM119 positive cells out of CD11b positive cells. Quantification was done with the 'sum area quantification' tool (n=4, each color represents a separate trial).
b) Slides were also stained with TMEM119 and IL33. Analysis was performed with Fiji ImageJ. The percentage of Tmem119 positive cells (microglia) out of the IL33 positive cells is presented.
c) The staining intensity of IL33 was measured for the TMEM119+ cells. Analysis was done with Fiji ImageJ (Fisher's Combined Probability Test; n=3, Within each trial, a Mann Whitney U test was performed to obtain the P-values, n=10-15 images per condition. One tailed).

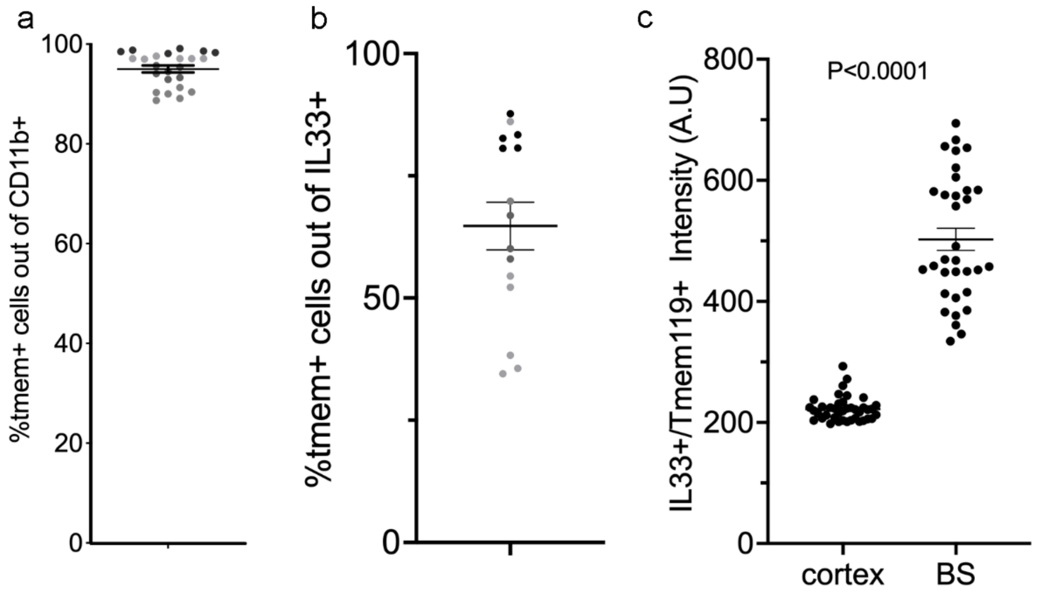


**Supplementary Figure 8: mouse IL33's anatomical differential expression is age dependent.** Three mice from each described age (P2, 2 months, 6 months, and 1 year) were sacrificed and RNA was extracted from their brainstems and cortices. The graph shows the relative higher expression of mIL33 in the brainstem in qPCR analysis. Each point is an average of three repeats. A significant decrease in higher BS expression is noted after the age of 2 months.

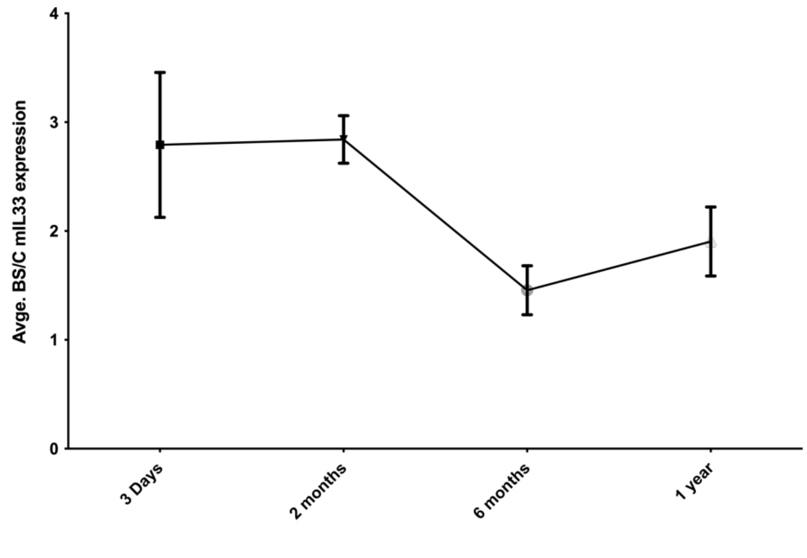


**Supplementary Figure 9: IL33 dose response**

About 1,000 SU-DIPGXIII cells were plated in 0.3 $cm^2$ wells, in about 80ul TSM based medium+factor/control. A serial dilution of the medium with recombinant IL33 was performed in order to prepare the appropriate concentrations, starting with 200ng/ml and diluting 1:1 in each round (all the way to 12.5ng/ml), and a control condition (0 ng/ml). For each concentration at least 7 wells were plated. After 72 hours of proliferation, the number of cells in each well was quantified by adding CellTiter-Glo reagent as previously described**.** The data presented is the average quantification measured for each concentration. A clear dose response was visible from 0 to 50 ng/ml, with maximal proliferation achieved at 50ng/ml, and so this concentration was chosen for the panel of experiments carried out (n=3).

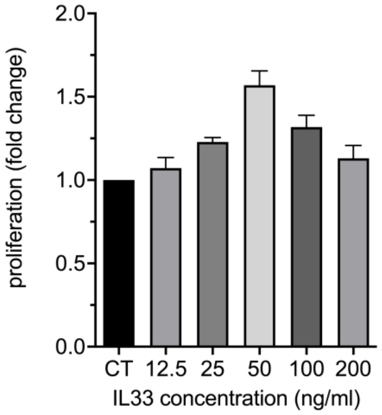


**Supplementary Figure 10:** Images of representative DIPG cells in 3D growth assay. SU-DIPGXIII cells were plated in high concentration in one spot (with either no addition or with recombinant IL33), covered with TSM:Matrigel (1:1) and grown for 7 days. At the end of the experiment, the cells were stained with crystal violet. The image demonstrates the DIPG cells' ability to grow/disseminate in a differential manner. There is a visibly noticeable increase in the cells' ability to grow with IL33 in comparison to the control condition. Scale bar: 100µm.

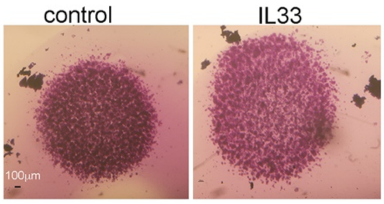


**Supplementary Figure 11:** The effect of recombinant IL33 on the average colony size per well was tested. Cells were plated as described in the colony count trial, and the colony size was measured with Incucyte® S3 Live-Cell Analysis system. The average colony size was higher in all DIPGs, though only 2 are significantly higher. (Fisher's Combined Probability Test; SU-DIPGXIII n=11, SU-DIPGXXV n=8, BT869 n=6. Within each trial, a Mann Whitney U test was performed to obtain the P-values, n=6-8 wells. One tailed). For each trial, the passage number is approximately indicated (P1-6; it was difficult to portray all of the repeats, so only the obvious clusters are marked).

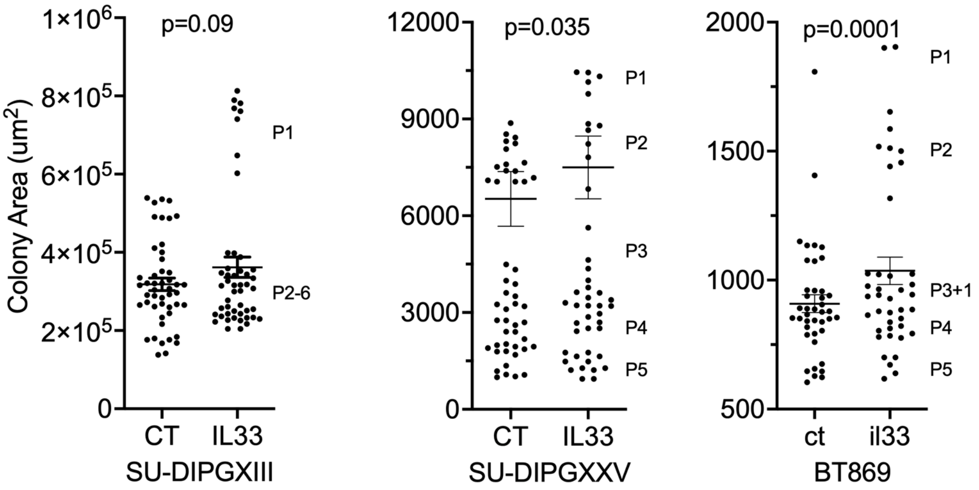


**Supplementary Figure 12: shRNA knockdown effect, SU-DIPGXIII**

qPCR results of knockdown of endogenous IL1RL1 in SU-DIPGXIII cells by two different shRNA sequences, as well as a CT sequence. The graph shows fold change in gene expression ±SEM as compared to the CT (data are presented as mean ± s.e.m., Mann–Whitney U test, n=5).

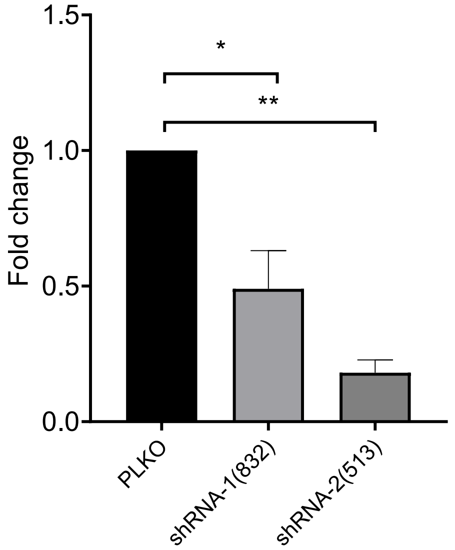


**Supplementary table1: Cortical microglia/brainstem microglia DEGs and their sub-cellular annotations.** Subcellular localization using Uniprot database. Ensembl identifiers and gene names were used as search input. Only annotations from reviewed entries (UniProtKB/Swiss-Prot) were included, and the two lists, resulting from Ensembl ID search (genes annotated only based on Ensembl ID marked in green font) and gene name (gene name only marked in red font) search respectively, were combined following annotation.
